# A KETOGENIC DIET ACCELERATES BLOOD-SPINAL CORD BARRIER RESEALING AND DECREASES SPINAL CORD IMMUNE CELL ACCUMULATION IN A PRECLINICAL MODEL OF MULTIPLE SCLEROSIS

**DOI:** 10.64898/2026.02.06.704406

**Authors:** Kendra S. Plafker, Dorothy A. Walton, Nathan Pezant, Susan Kovats, Robert C. Axtell, Scott M. Plafker

**Affiliations:** Aging and Metabolism Research Program, Oklahoma Medical Research Foundation, Oklahoma City, OK USA; Center for Biomedical Data Sciences, Oklahoma Medical Research Foundation, Oklahoma City, OK USA; Arthritis and Clinical Immunology Research, Oklahoma Medical Research Foundation, Oklahoma City, OK USA; Department of Cell Biology, University of Oklahoma Health Sciences Center, Oklahoma City, OK USA

**Keywords:** Ketogenic diet, multiple sclerosis, EAE, blood-CNS barrier, immune cell

## Abstract

We previously reported that a medium chain triglyceride and *ω*-3 fatty acid enriched ketogenic diet (KD) restored vision and motor functions to mice undergoing experimental autoimmune encephalomyelitis (EAE), a preclinical model of multiple sclerosis (MS). In the mechanistic studies reported here we show that the KD dramatically reduces the infiltration of CD45**^HI^** immune cells into the spinal cords of EAE mice when initiated at symptom onset and fed for 2 weeks. Two mechanisms contributed to this reduced accumulation of infiltrates. First, the KD accelerated resealing of the blood-SC barrier (BSCB) in both sexes. This resealing accompanied motor score functional recovery and occurred whether KD feeding was initiated the day of symptom onset or 1 week after symptom onset. Consistent with this resealing, the KD reduced SC levels of soluble IL-1β, a primary cytokine associated with compromising the integrity of CNS-blood barriers. A second mechanism was that the KD increased the percentage of spinal cord infiltrates undergoing apoptosis. Beyond reducing the total abundances of immune cell infiltrates by these two mechanisms, the KD also preserved subpopulations of neutrophils and non-classical mono/macs expressing surface markers associated with inflammation resolution, healing, and tissue repair. Studies leveraging global knockout mouse strains showed unexpectedly that the efficacy of the KD in this interventional paradigm was independent of both the hydroxycarboxylic acid receptor 2 (HCA_2_) and the anti-stress transcription factor, Nrf2. Collectively, this work advances novel insights by which a KD ameliorates autoimmune-mediated MS-like pathologies to promote functional recovery and more broadly, implies that this dietary strategy has potential benefit for facilitating the healing of disrupted blood-CNS barriers and resolving inflammation in multiple neurodegenerative diseases.

## INTRODUCTION

MS is the most common neurodegenerative disease in young adults with nearly 3 million individuals afflicted worldwide (1). Experimental autoimmune encephalomyelitis (EAE) rodent models reconstitute some hallmark pathologies of MS (2) and have been instrumental in developing disease modifying therapies (DMTs) for treating people with MS (pwMS) (3). Similar to other autoimmune disorders, MS is more prevalent in females compared to males, with a ≥ 2-fold greater incidence among females (4). This female propensity has resulted in most EAE studies being performed in female mice (5), yet males with MS can have more severe disability (e.g., (6)), underscoring the need to investigate both sexes using the EAE paradigm to identify common as well as sex-divergent responses to candidate interventions. Further, the immune systems of females and males differ in humans and mice, and EAE studies specifically designed to identify how the model differs between the sexes have revealed both common and differential immune cell responses (5, 7). It is postulated that such differences contribute to the varied DMT efficacies, preferences, and adherences to treatment regimens between sexes (e.g., (8, 9)), but to date, sufficiently powered studies have not identified sex-based differences in DMT efficacies in pwMS.

Ketogenic diets (KD) are high-fat, very low-carbohydrate dietary regimens that induce a metabolic state of ketosis, in which ketone bodies, including beta-hydroxybutyrate (β−HB), become the primary substrates for energy production in place of glucose. KDs were originally developed in the 1920s as a therapeutic intervention for pediatric drug-resistant epilepsy and have since been investigated for potential applications in weight management, metabolic disorders, neurological diseases, and other clinical conditions (10).

KD and carbohydrate restriction studies in rodent EAE models have demonstrated improvements in motor functions and vision as well as spatial learning and memory (11–15). These improvements were accompanied by reductions of pro-inflammatory cytokines and chemokines (11–14). We previously reported that a non-obesogenic KD enriched in fiber, medium chain triglycerides (MCTs; octanoic acid and decanoic acid), and ω-3 fatty acids prevented symptom onset when fed prophylactically to female and male EAE mice. The diet likewise restored visual and motor function to both sexes when initiated as an intervention, after symptom onset (11). To identify the mechanisms underlying KD efficacy, we performed immune cell profiling studies of the spinal cords (SCs), blood, and bone marrow (BM) of female and male mice undergoing EAE. For all experiments, the KD was initiated as an intervention after symptom onset to reflect the timing by which this dietary strategy would likely be leveraged by pwMS in response to symptom flare-ups or by individuals with clinically isolated syndrome that have not received a definitive MS diagnosis and are DMT-naïve.

We report here that both female and male mice had dramatic reductions of total CD45**^HI^** infiltrating immune cells in the SC after being fed the KD for 2 weeks with neutrophils being reduced the most (i.e., > 90%). Preceding this reduction, neutrophils were elevated in the circulation within a week of diet initiation. Mechanistically, these altered immune cell distributions in the SC and blood indicated that the KD accelerated resealing of the blood-spinal cord barrier (BSCB), which was confirmed experimentally using a fluorescent tracer. Remarkably, this resealing was also observed when the diet was initiated a week after symptom onset. The accelerated BSCB resealing corresponded with a reduction of soluble IL-1β levels in the CNS of both sexes and in females, was also associated with increased expression of adherens-and tight-junction proteins, VE-cadherin and ZO-1, respectively. We also discovered that the KD reduced the accumulation of pathogenic immune cells by promoting the apoptosis of multiple adaptive and innate immune cell types within the spinal cord while preserving SC-infiltrated neutrophils and monocyte/macrophages bearing markers associated with anti-inflammatory, pro-resolution phenotypes. Lastly, we found that the efficacy of the KD in this interventional paradigm was independent of the hydroxycarboxylic acid receptor 2 (HCA**_2_**) as well as the anti-stress transcription factor, Nrf2.

## MATERIALS AND METHODS

### EAE procedure and mice

7.5-13 week old mice were immunized with 250 ug of a peptide corresponding to amino acids 35-55 of murine MOG emulsified in incomplete Freund’s adjuvant supplemented with 250 ug heat-inactivated *Mycobacterium tuberculosis* (Thermo Fisher Scientific; DF3114338). 250 ng of *Bordetella pertussis* toxin was administered the day of immunization and again 2 days later. Clinical symptoms began 11-16 days post-immunization. Mice were weighed daily and scored for motor deficits daily using our published scoring system (11). Wild type C57BL/6J mice were used for experiments involving immunohistochemistry, sodium fluorescein and ELISAs. A transgenic C57BL/6J strain we call ‘SMART-Fox’ was used for flow cytometry experiments to circumvent the need for intracellular staining. SMART-Fox mice were generated by crossing Smart-17A mice (JAX stock # 032406;) to B6-Foxp3^EGFP^ mice (JAX stock # 006772) and are homozygous for each reporter construct. Males are hemizygous for the Foxp3-GFP transgene because this construct is on the X chromosome. SMART-17A mice express a non-signaling form of the human nerve growth factor receptor upon activation of the IL-17A locus. The B6-Foxp3^EGFP^ mice co-express enhanced GFP and Foxp3 under the endogenous *Foxp3* promoter with GFP expression accurately marking the Foxp3^+^ CD4^+^ Treg population. Notably, the two parental strains have been validated in the MOG**_35-55_**-EAE model (16, 17). Importantly, we observe similar motor deficits in SMART-Fox mice and wild type C57BL/6J mice. Blood ketones were measured using Precision Xtra Blood Ketone Monitoring System.

HCA**_2_** knockout mice on a C57BL6/J background were obtained from Dr. Muthusamy Thangaraju (University of Augusta, GA) after obtaining permission from the investigator that created this global knockout strain, Dr. Stefan Offermanns (Max Planck Institute for Heart and Lung Research, Germany). Nrf2 knockout mice were purchased from Jackson Labs (cat # 017009) and backcrossed onto a C57BL/6J background to remove (as confirmed by PCR) the retinal degenerative RD8 mutation.

### Diet compositions

Teklad control (TD.170645) and ketogenic (TD.10911) diets were obtained from Envigo, Inc. The KD provides 4.7 Kcal/g with 22.4% Kcal from protein, 0.5% Kcal from carbohydrate, and 77.1% Kcal from fat. The control diet (CD) provides 3.6 Kcal/g with 20.4% Kcal from protein, 69.3% Kcal from carbohydrate, and 10.4% Kcal from fat. Diet ingredients are detailed in our previous publication (11). Food and water (containing 2.6 ml/L electrolytes, Electrolyte drops; KetoChow) were provided *ad libitum* and additional hydration/electrolytes were provided using intraperitoneal saline injections to animals showing signs of dehydration.

### Intervention Diet studies

Unless otherwise indicated, all studies in this manuscript used an intervention paradigm in which the animals are maintained on standard chow until being switched to either the CD or KD after motor symptom onset and then maintained on diet for the indicated times.

### Therapeutic Diet study

In the therapeutic paradigm used exclusively for Fig. 6, mice were maintained on standard chow until 6-8 days post motor symptom onset. At that time, only mice that had reached a motor score of at least 2.5 within the first 6-8 days were switched to CD or KD for 2 additional weeks before harvest.

### Multiparameter Flow cytometry

At the indicated times, animals were sacrificed by CO_2_ asphyxiation, blood was collected by cardiac puncture followed by perfusion with PBS and decapitation. RBCs were lysed in 100ul blood, remaining cells were washed and stained. Bone marrow was flushed from one leg (femur + tibia) per mouse, RBCs were lysed and 5 million cells were stained. Spinal cords were dissected, dissociated mechanically and incubated in collagenase and DNase. Cells were filtered through a 70uM mesh filter, washed with PBS and myelin was removed using a 40% Percoll gradient; all remaining spinal cord cells were stained. Cells were incubated with BD Horizon Fixable Viability Stain 450 and anti-CD16/32 for Fc block. Supplemental Table 1 lists all the fluorophore-conjugated surface staining antibodies used. Cells were collected on a Cytek Aurora 5 laser instrument and analyzed using FlowJo v10. All raw flow data was processed sequentially as follows: 1) all debris was gated out, 2) singlets were gated by FSC-area and-height and SSC-area and-height and 3) lastly, dead (viability stain-positive) cells were gated out. Additional organ-specific gating is illustrated in Supplemental Figure 2.

### Immunohistochemistry (IHC)

Fixed, paraffin-embedded spinal cord sections were de-paraffinized, heated in antigen retrieval citrate buffer in a Retriever 2100 (Aptum Biologics, Inc.) and then incubated with anti-proteolipid protein (α-PLP) (rat: 1:200 dilution; Millipore, Inc.). Nuclei were counterstained with Hoechst, and images were captured using a Zeiss Axioscan 7 digital slide scanner housed in the OMRF Imaging Core.

### Sodium fluorescein assay

Healthy or EAE mice at either 0 days post symptom onset (DPSO), 3 DPSO, or 6 DPSO (Fig. 5) or alternatively at 21 DPSO (Fig 6) were intraperitoneally injected with 100ul of 10% sodium fluorescein (NaFl). 30 minutes post-injection, spinal cords and blood were collected to quantify extravasated fluorescent tracer and serum concentrations, respectively. NaFl was quantified using a SpectraMax M2 plate reader (Molecular Devices).

### ELISAs

Spinal cords were collected, snap frozen in liquid nitrogen and stored at-80°C to simultaneously measure cytokines from all timepoints and experiments. Spinal cords were weighed, thawed and mechanically dissociated in PBS containing EZBlock protease inhibitor cocktail (Biovision). The analytes measured were IL-6 (Invitrogen, cat#5017218), IL-1β (Invitrogen, cat# 887013A22), and CXCL2 (R&D, MM200) using a SpectraMax M2 plate reader (Molecular Devices).

### Apoptosis Assay/FLICA

For the FLICA/AnnexinV assay, live cells were incubated with YVAD-FLICA660 (Immunochemistry Technologies, Cat No. 9122; diluted 1:125) at 37°C for 30 min. After staining for viability and surface markers as described above, cells were stained with Andy Fluor™ 555 Annexin V (ABP Biosciences) per the manufacturer’s instructions. Cells were fixed and permeabilized for intracellular IL-1β using BD Cytofix/Cytoperm™ Kit (Cat. No. 554714).

### Western blotting

Spinal cords of healthy mice and EAE mice at Peak disease were collected in T-PER (Thermo Scientific, Cat. No. 78510) following manufacturer’s instructions. 100ug protein from each SC was subjected to Western blot analysis as previously described (18). Antibodies used were anti-ZO-1 (ABCAM, Cat. No. AB190085), anti-VE Cadherin (ABCAM, Cat. No. AB33168) and HRP-conjugated anti-β-Actin (Proteintech, Cat. No. HRP-60008).

### Statistical and T-distributed stochastic neighbor embedding (tSNE) Analyses

Graphs show average of all data points per sex and treatment with error bars indicating standard error of the mean. Two-tailed, unpaired Student’s t test was used unless a non-normal distribution was seen by Shapiro-Wilk test in which case, Mann-Whitney U test was used. tSNE was performed in FlowJo using all fluorescent markers except viability. Auto (opt-SNE) learning configuration (1000 iterations, perplexity=30) using Approximate (random projection forest-ANNOY) KNN algorithm and FFT Interpolation (Flt-SNE) gradient algorithm were used.

## RESULTS

Our previous work showed that a KD restored visual and motor function to mice undergoing the MOG**_35-55_** EAE mouse model of autoimmune-mediated inflammation and demyelination (11). The present studies extend these findings by applying immune cell profiling to identify mechanistic insights by which the KD mediates therapeutic benefits. As we previously observed in wild type C57BL/6J mice (11) and repeated here in a reporter strain called ‘SMART-Fox’ (C57BL/6J background), switching EAE mice to the KD following symptom onset resulted in functional motor recovery beginning within 2 days for females and within 2-4 days for males. This rapid recovery persisted until the termination of experiments at 14 days DPSO (i.e., equal to 14 days of diet treatment) (Figs. 1A and 1B, respectively). As further detailed in the Material and Methods section, SMART-Fox mice express a reporter for IL-17A expression as well as enhanced GFP under the endogenous *Foxp3* promoter. Notably, only the GFP reporter was tracked in the present study. KD-mediated motor function recovery corresponded with increased ketones in the circulation (Fig. 1C), nominal variations in body weight (Fig. 1D), and reduced SC lesion sizes and cellularity observed by immunohistochemistry (IHC) (Supplemental Fig. 1).

**Figure 1.**
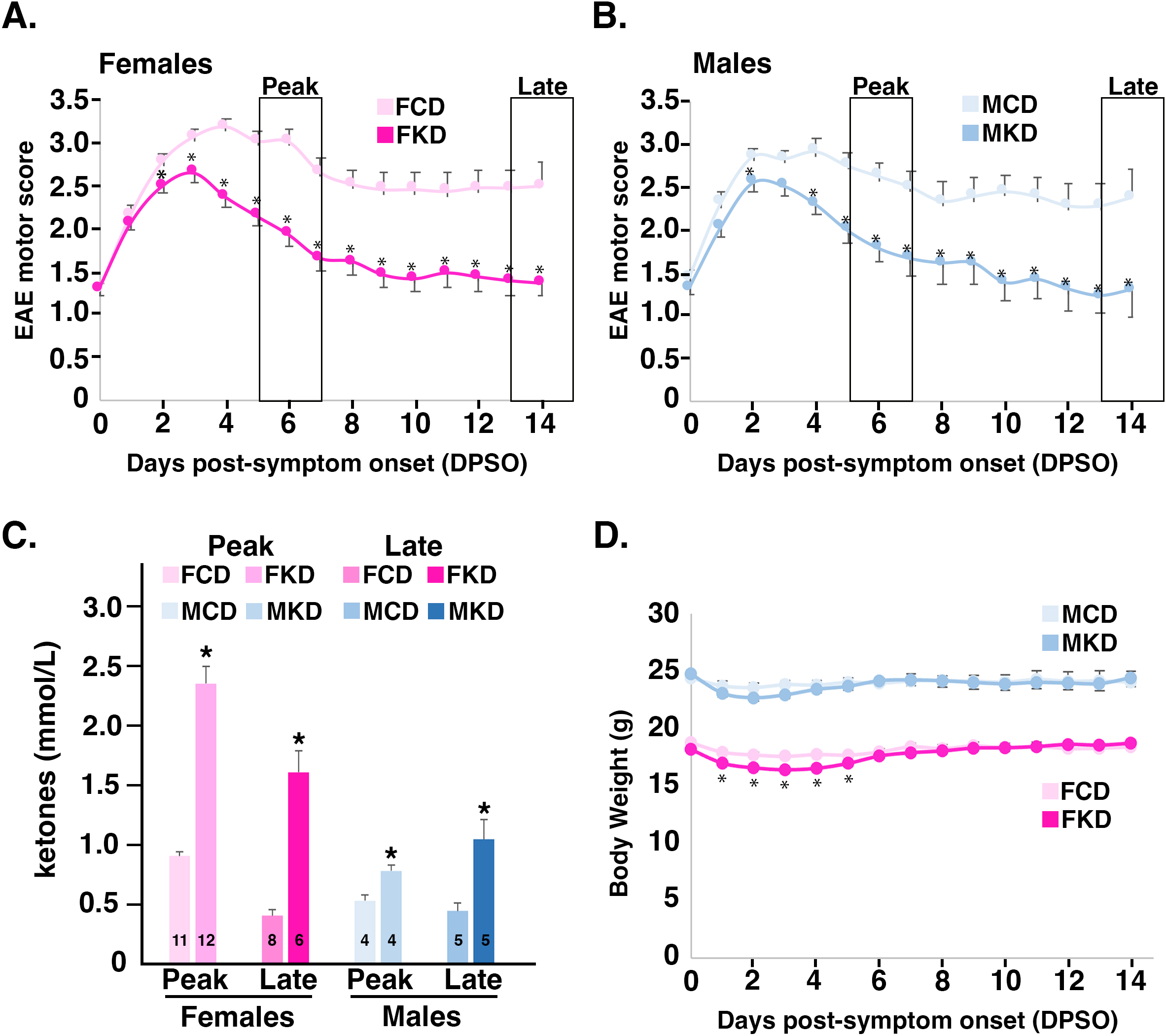
The KD restores motor functions to female and male mice undergoing EAE. SMART-Fox mice immunized with MOG_35-55_ were fed either KD or CD following symptom onset (motor score ≥ 1) and maintained on their respective diets for 14 days. Motor scores were tracked daily for females (**A**) and males (**B**). n=15-17 mice/sex/diet/time point. ‘Peak’ disease represents 5-7 days post-symptom onset (DPSO) and ‘Late’ disease represents 13-15 DPSO, the two time points when tissues were harvested for most of the subsequent figures. (**C**) Blood ketone levels (mmol/L) measured at Peak and Late disease. The number of mice used for measurements is indicated on the data bars. (**D**) Body weights of mice in grams during the period starting from the start of KD or CD treatment until the end of the experiments. Sexes and diets are indicated using the following abbreviations: EAE female mice on CD (FCD), EAE female mice on KD (FKD), EAE male mice on CD (MCD), and EAE male mice on KD (MKD). These abbreviations are used throughout for labeling figures. Asterisks denote statistical significance (p<0.05) as determined by two-tailed, unpaired Student’s t test.

To circumvent the inherent limitations and variability of quantifying and characterizing infiltrates in tissues using IHC (19, 20), we used multiparameter flow cytometry with validated fluorochrome-labeled monoclonal antibodies (Supplemental Table 1) to profile the immune cell changes mediated by the KD in the SC (Fig. 2), blood (Fig. 3), and BM (Fig. 4). Profiling was done at two time points, 5-7 DPSO (aka ‘Peak’ disease) and 13-15 DPSO (aka’Late’ disease) (Fig. 1). The surface markers used to define each immune cell type and the abbreviations for all immune cells are compiled in Supplemental Table 2; the flow cytometry gating parameters are provided in Supplemental Fig. 2. All flow cytometry experiments were performed in the SMART-Fox strain to facilitate visualizing FoxP3**^+^** T regulatory cells (Tregs) without the need for intracellular transcription factor labeling.

**Figure 2.**
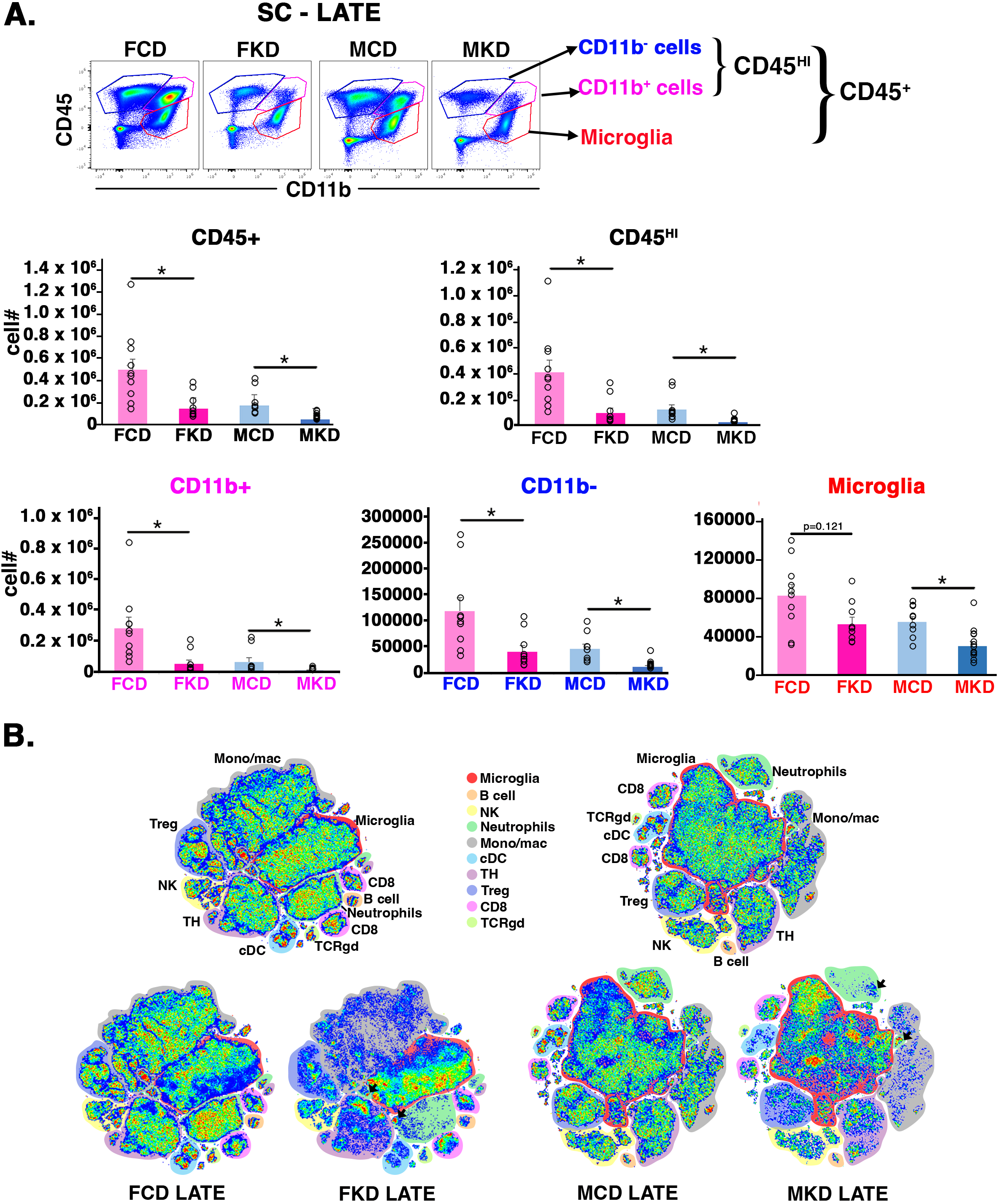
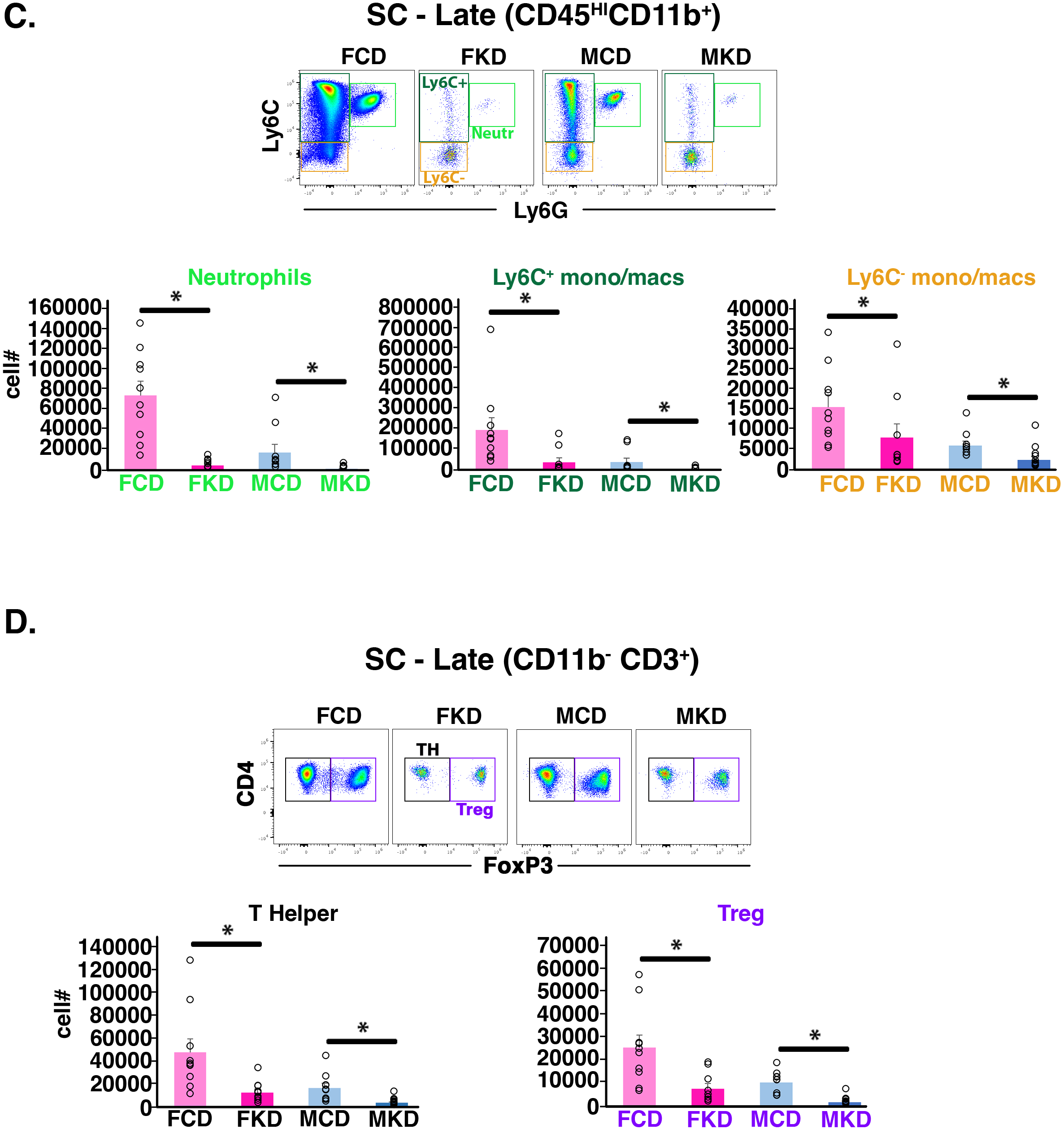
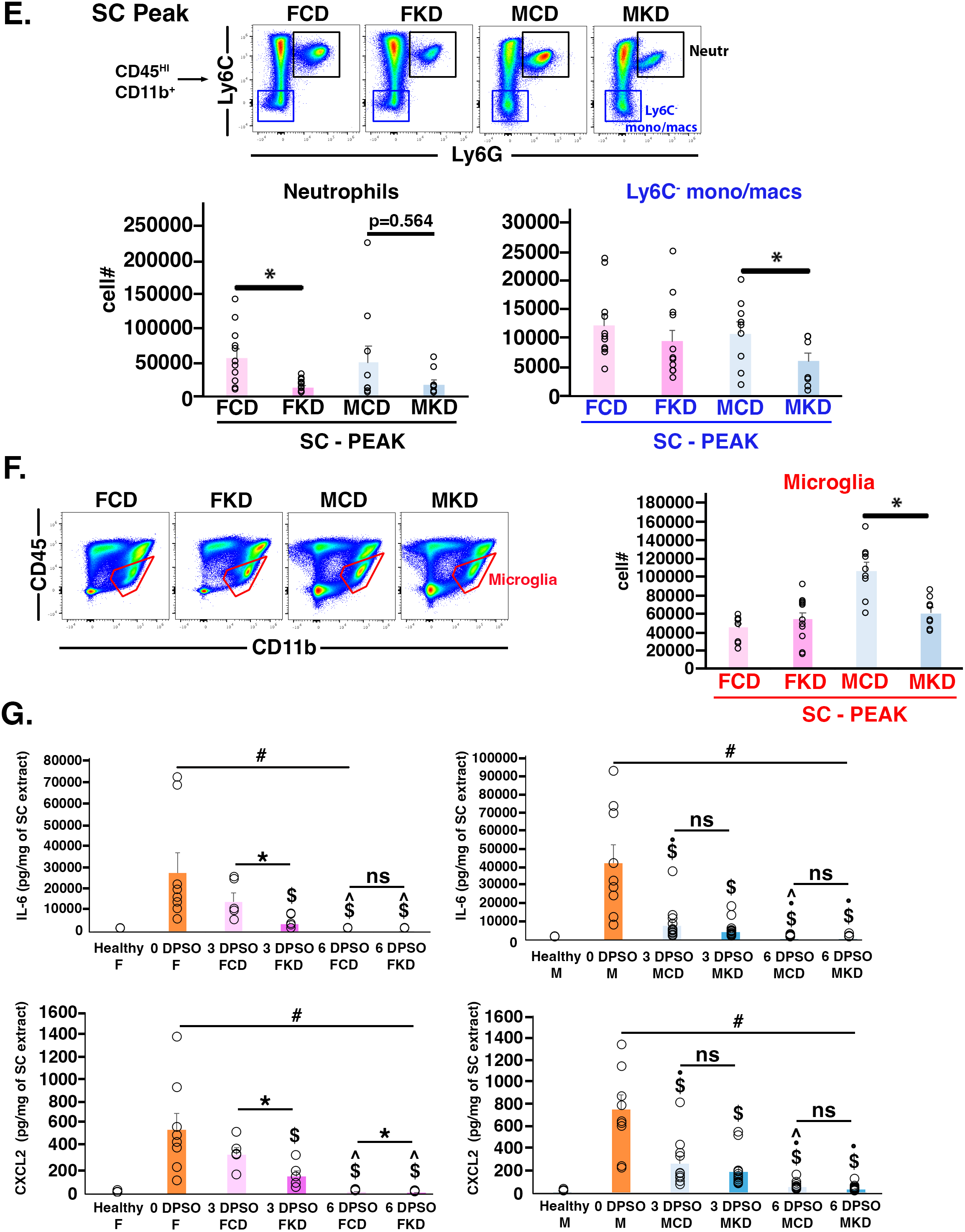
The KD reduces cell types that mediate EAE pathogenicity in the SCs of female and male mice. **(A)** Representative flow cytometry pseudo-colored density plots of live SC immune cells at Late disease gated on CD45 and CD11b expression. CD45**^HI^** cells are comprised of CD45**^HI^** CD11b**^-^** and CD45**^HI^** CD11b**^+^** cells, as marked in blue and pink polygons, respectively. Total CD45**^+^** cells are comprised of CD45**^HI^** cells plus microglia (marked in the red polygon). Graphs show abundances of the indicated cell types for each animal (open circles) as a function of diet and sex. (**B**) t-SNE plots generated from flow cytometry data of Late Disease SC immune cell profiling. Female plots are to the left of the key and male plots are to the right. Top plots (adjacent to key) are composites of CD and KD plots. Below each composite is the plot for CD on the left and KD on the right. Cell populations are color coded in all plots as defined in the key. Black arrows on FKD Late plot and MKD Late plot mark neutrophil populations retained in KD fed mice. Data for t-SNE analysis from n=3 mice/sex/diet. (**C,D**) Representative flow cytometry plots generated from live SC CD45**^HI^** CD11b**^+^** cells isolated at the Late time point. In (**C**), cells gated on Ly6C and Ly6G expression giving rise to graphs of neutrophils (lime green box), Ly6C**^+^** mono/macs (dark green box), and Ly6C**^-^** mono/macs (orange box) abundances. In (**D**), live SC CD45**^HI^** CD11b**^-^**CD3^+^ TCRb^+^ T cells gated on CD4 and FoxP3 expression (i.e., GFP) giving rise to graphs of TH cell (black box) and Treg (purple box) abundances as a function of diet. (**E**) Representative flow cytometry pseudo-colored density plots gated on Ly6C and Ly6G expression of live CD45**^HI^**CD11b**^+^** immune cells recovered from SCs at Peak disease giving rise to graphs of neutrophils (black box) and Ly6C**^-^** mono/macs (blue box). (**F**) Representative flow cytometry pseudo-colored density plots of live immune cells recovered from SCs at Peak disease with gating on CD45 and CD11b expression to label microglia, marked in red polygons. The accompanying graph shows average microglial cell number and data from individual animals. In (A-F), asterisks denote statistical significance * p<0.05 as determined by Mann-Whitney U test. All graphs were constructed from flow cytometry data, with all cell types profiled shown in Supplemental Table 3. n=9-13 mice/sex/diet. (**G**) Graphs of lL-6 and CXCL2 quantified from SC extracts as a function of sex, diet, and DPSO (equivalent to the number of days mice were fed KD or CD after symptom onset). Left graphs are female data, and right graphs are male data. n=5-13 mice/sex/treatment. For (**G**), asterisks denote statistical significance **\*** p<0.05 vs CD; # indicates p<0.05 vs Healthy; **$** indicates p<0.05 vs 0 DPSO; **^** indicates p<0.05 vs 3 DPSO; and • indicates p<0.05 vs female as determined by Mann-Whitney U test.

**Figure 3.**
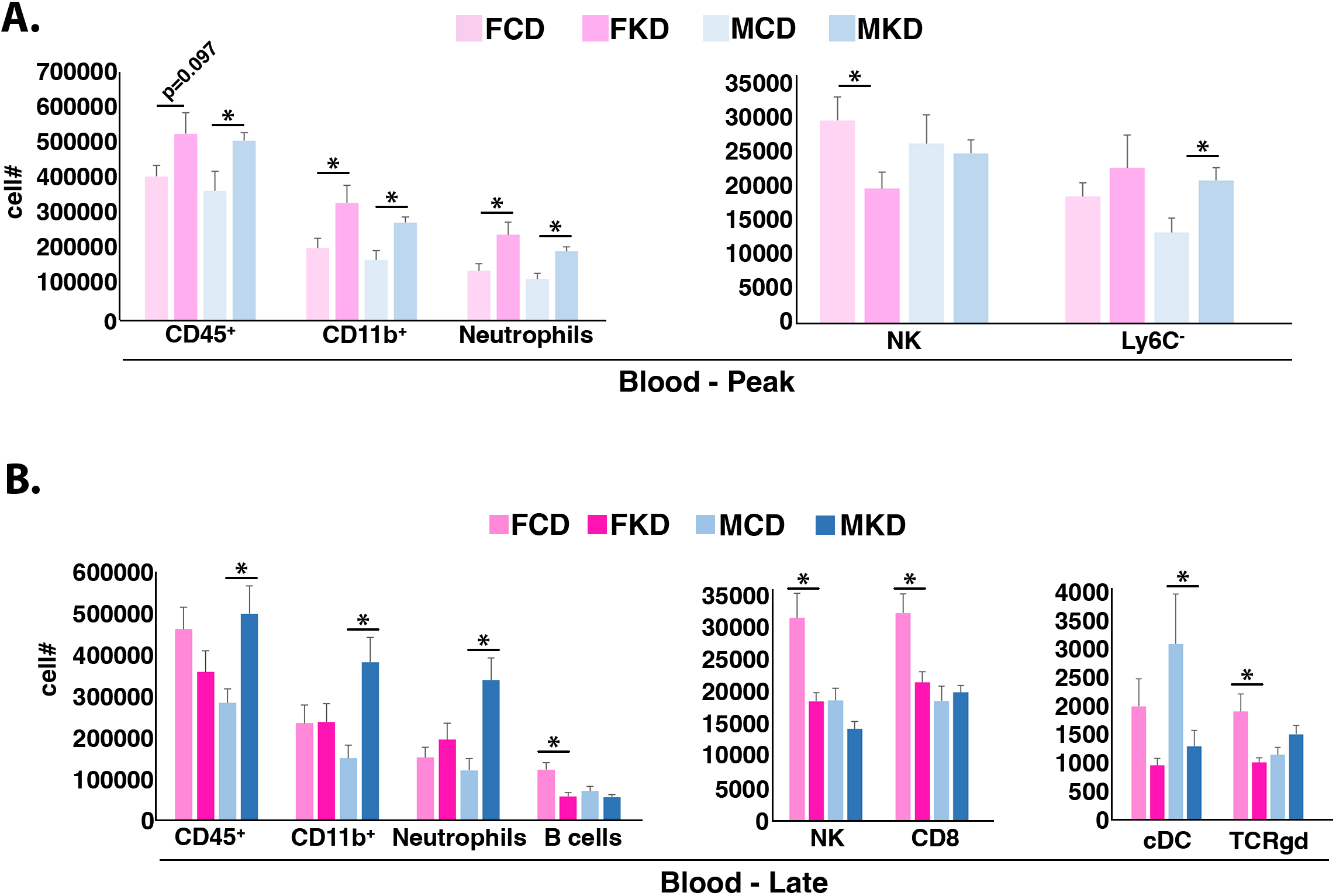
The KD increases neutrophils in the circulation. (A,B) Graphs depicting the abundances of different cell types that changed at Peak disease (**A**) or Late disease (**B**) in the blood as a function of sex and diet. Graphs were constructed from flow cytometry data, and all cell types profiled (including those that did not change) are provided in Supplemental Table 3. n=9-13 mice/sex/diet. Asterisks denote statistical significance * p<0.05 as determined by two-tailed, unpaired Student’s t test.

**Figure 4.**
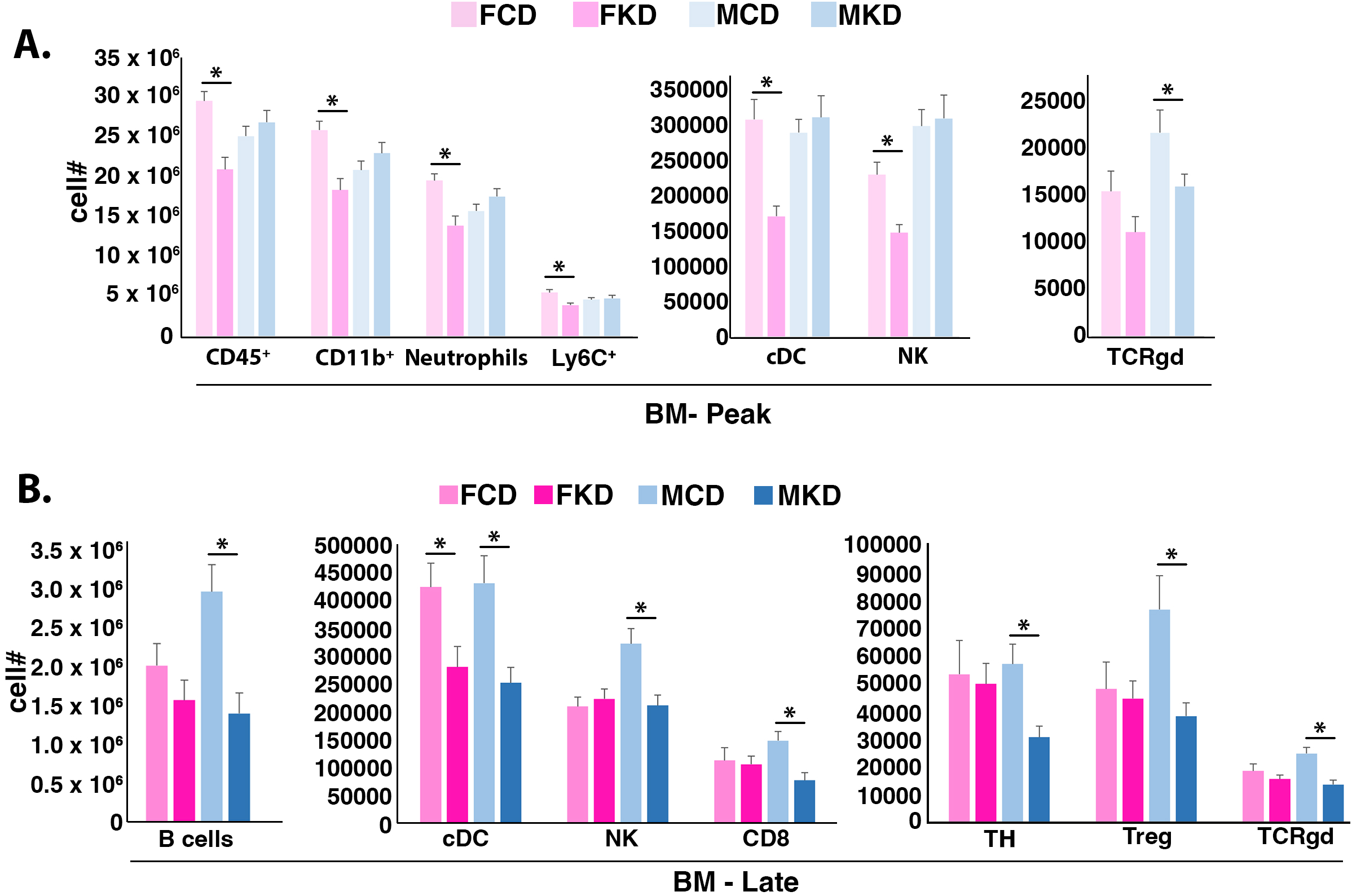
The KD decreases multiple immune cell populations in the bone marrow (BM). (A, **B)** Graphs depicting the abundance of cell types that changed at Peak disease (**A**) or Late disease (**B**) in the BM as a function of sex and diet. All graphs were constructed from flow cytometry data, with all cell type raw data shown in Supplemental Table 3. n=9-13 mice/sex/diet. Asterisks denote statistical significance * p<0.05 as determined by two-tailed, unpaired Student’s t test.

**Figure 5.**
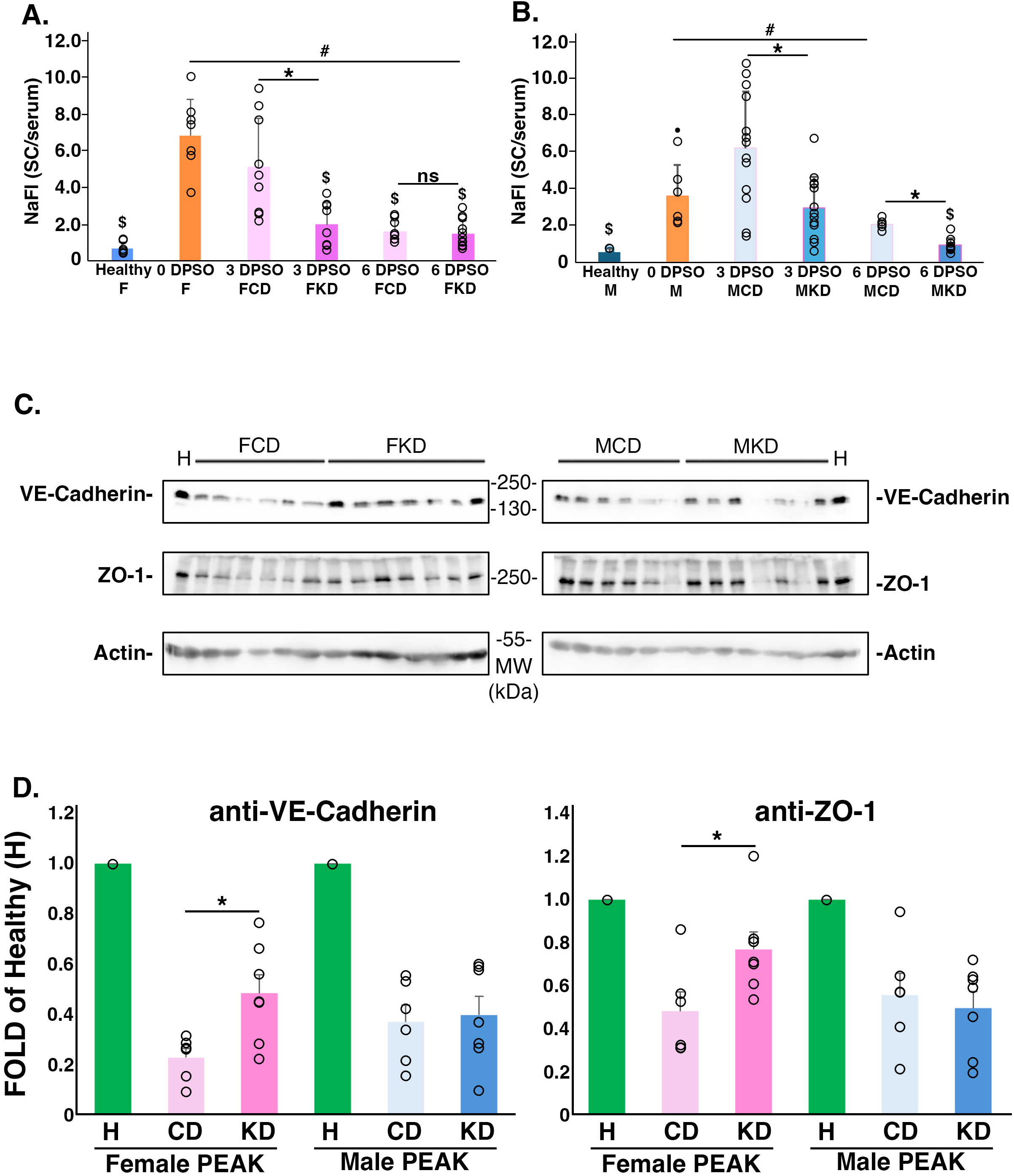
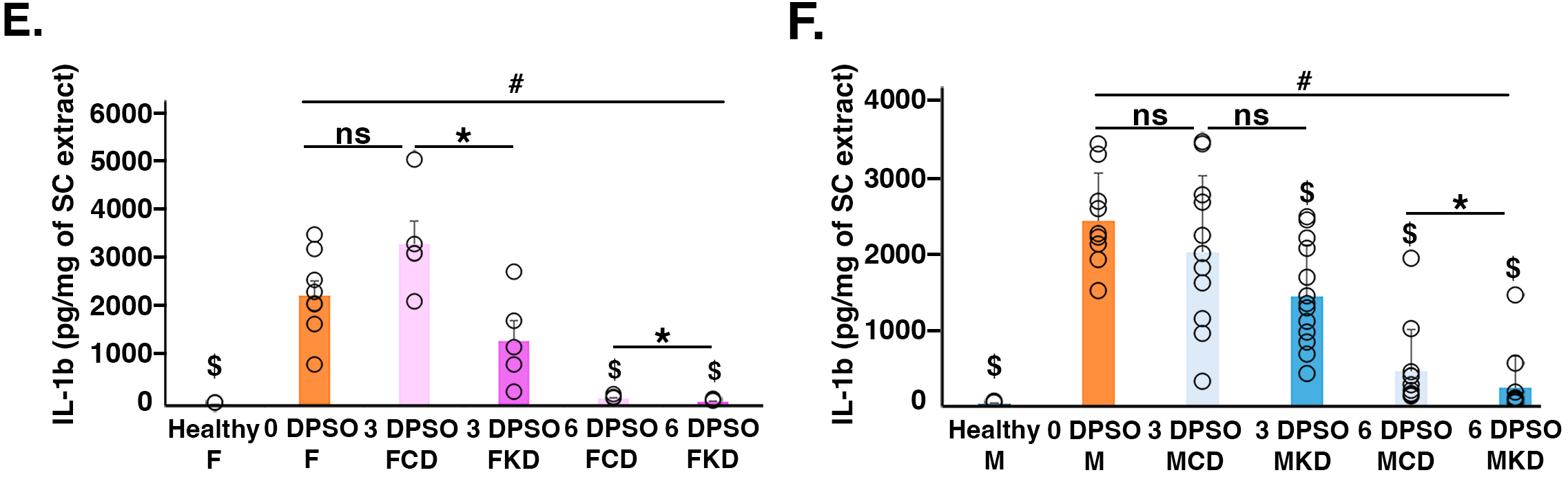
The KD accelerates resealing of the BSCB. (A,B) Graphs of extravasated sodium fluorescein (NaFl) isolated from SC extracts at the indicated times DPSO. NaFl is expressed on the y-axis as the ratio of the fluorescent tracer in the SC relative to the amount in the serum. Each open circle represents an individual animal. “Healthy” are age-and sex-matched non-EAE mice eating standard chow. n=4-13 mice/sex/diet. (**C**) The KD increases VE-cadherin and ZO-1 expression in females. SCs were recovered from female and male EAE mice after 5-7 days of treatment with the KD or CD. SC lysates were probed with the indicated antibodies to VE-cadherin, ZO-1, and actin. The migrations of molecular weight markers are indicated adjacent to each blot. ‘H’ – SC extract derived from a healthy (non-EAE), strain-, age-, and sex-matched mouse. (**D**) Graphs of quantified Western blot bands. Band intensities indicated as fold change compared to VE-cadherin (left graph) and ZO-1 (right graph) expression levels in healthy mice. Open circles indicate individual band measurements, and asterisks denote statistical significance as determined by unpaired, two-tailed Student’s t test. (**E,F**) Graphs of ELISA data quantifying soluble lL-1β in SC extracts as a function of sex, diet, and DPSO (equivalent to the number of days mice fed KD or CD). Soluble IL-1β was recovered from mice at 0 DPSO, 3 DPSO, and 6 DPSO. n=5-13 mice/sex/diet. For (**A,B,E,F**), asterisks denote statistical significance **\*** p<0.05 vs CD; # indicates p<0.05 vs Healthy; **$** indicates p<0.05 vs 0 DPSO; and • indicates p<0.05 vs Female as determined by Mann-Whitney U test.

**Figure 6.**
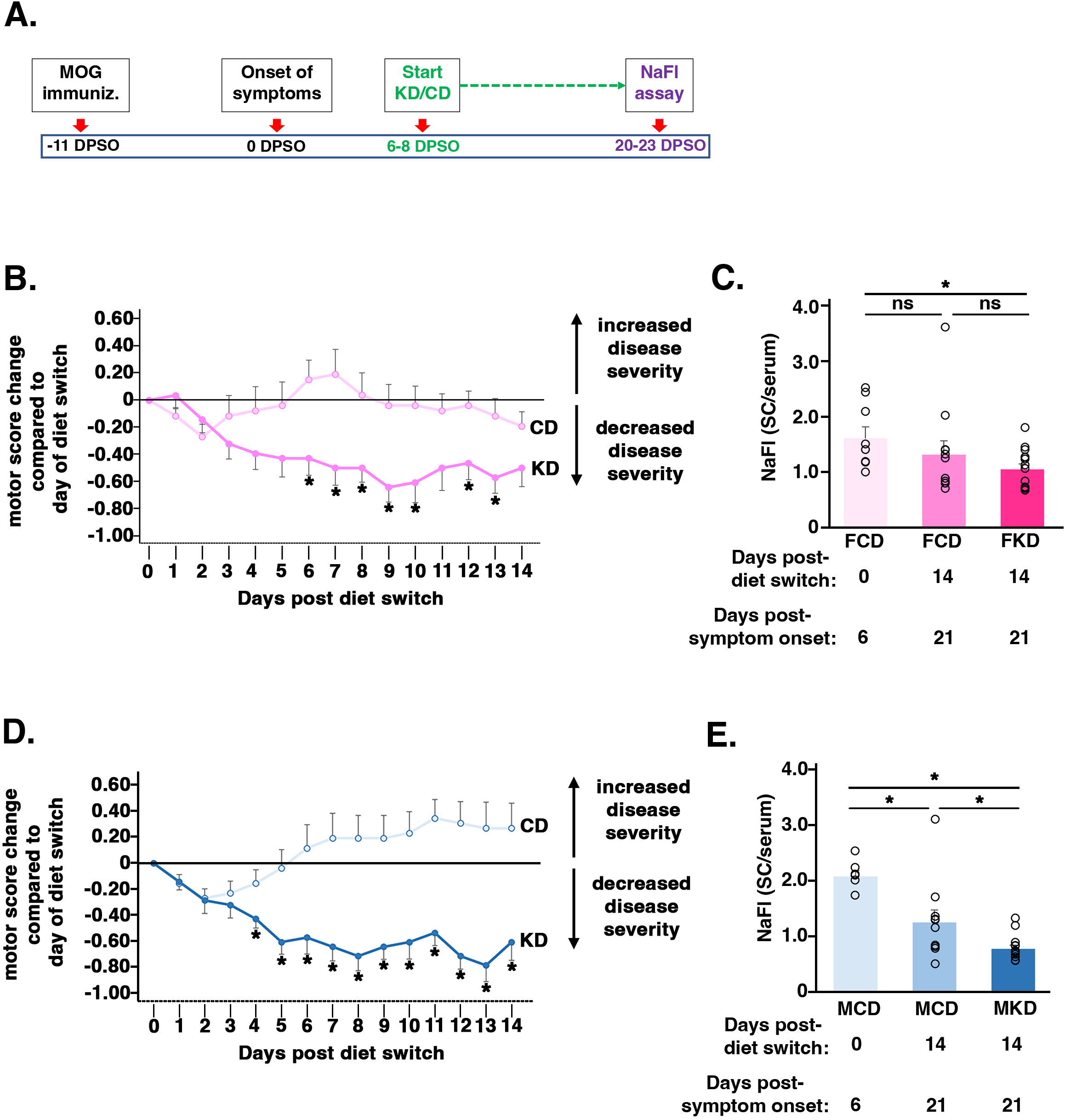
The KD improves motor function and reseals the BSCB when initiated 1 week after clinical symptom onset. **(A)** Schematic of study design to test if the KD promotes BSCB resealing and/or improves motor function when initiated 1 week after symptom onset. **(B,D)** Graphs of motor score changes each day compared to the motor score on the day of diet switch (i.e., 6-8 DPSO) for females (B) and males (D); n=10-13 mice/sex/diet. **(C,E)** Graphs of extravasated NaFl expressed as a ratio of the tracer in the SC relative to the serum for EAE females (C) and EAE males (E). Open circles represent individual animals. n=10-13 mice/sex/diet. 6 DPSO data derived from Fig. 5A and 5B is provided for comparison to highlight the capacity of the KD to further reduce BSCB permeability beyond the spontaneous resealing observed at 6 DPSO. Asterisks denote statistical significance (p <0.05) and ns = not statistically significant.

### The KD reduces immune cells in the SC of EAE mice

The major finding in the SCs of both sexes was that the KD drastically reduced the abundance of total CD45**^+^** immune cells by Late disease (Fig. 2A and Table 1), consistent with the observed motor function recovery (Fig. 1). The abundances of infiltrating CD45**^HI^** cells, including CD11b**^+^** myeloid populations and CD11b**^-^** cells (i.e., T cells, B cells, cDCs) were likewise reduced (Fig. 2A). Visualization of the data by t-SNE indicated that the KD reduced neutrophils, monocyte/macrophages (mono/macs), T helper cells (TH), and Tregs in the SCs of both sexes (Fig. 2B). These changes in cell abundances were confirmed to be statistically significant (Figs. 2C-D). By contrast, at the earlier time point (i.e., Peak), only neutrophils were decreased in female SCs and CD45**^HI^** CD11b**^+^**Ly6C^-^ mono/macs and microglia were decreased in male SCs (Figs. 2E and 2F, respectively and Table 1). Consistent with the early neutrophil reductions in female SCs, ELISA assays of SC extracts at 0, 3, and 6 days post-symptom onset (DPSO) showed that in females the KD decreased levels of the neutrophil activator, IL-6, at 3 DPSO whereas no statistically significant changes were detected in males (Fig. 2G). Likewise, CXCL2, the neutrophil recruitment cytokine, was reduced by the KD in females at both 3 and 6 DPSO but not in males (Fig. 2G).

**Table 1.** Immune cell profiling of SCs, blood, and BM. Tissues were harvested from EAE mice at Peak or Late disease. Immune cell profiling was done by multiparameter flow cytometry using validated fluorochrome-labeled monoclonal antibodies. Tables list fold change of KD compared to CD. Cell types that reached statistical significance are assigned fold change and blank cells indicate that statistical significance was not reached for that cell type. Statistically significant differences were determined by two-tailed, unpaired Student’s t test for blood and BM and by Mann-Whitney U test for SC. Heatmaps show **green** entries representing fold decreased cell abundance for KD and red entries representing fold increased cell abundance for KD. The raw data to generate this table are provided in Supplemental Table 3.

|  | Peak |  |  |  |  |  | Late |  |  |  |  |  | KD/CD |
| --- | --- | --- | --- | --- | --- | --- | --- | --- | --- | --- | --- | --- | --- |
|  | FOLD CHNG v CD* |  | FOLD CHNG v CD** |  | FOLD CHNG v CD** |  | FOLD CHNG v CD* |  | FOLD CHNG v CD** |  | FOLD CHNG v CD** |  | FOLD CHNG |
|  | SC | SC | Blood | Blood | BM | BM | SC | SC | Blood | Blood | BM | BM |  |
|  | PEAK | PEAK | PEAK | PEAK | PEAK | PEAK | LATE | LATE | LATE | LATE | LATE | LATE | 3.00 |
|  | FKD | MKD | FKD | MKD | FKD | MKD | FKD | MKD | FKD | MKD | FKD | MKD | 2.00 |
|  |  |  |  |  |  |  |  |  |  |  |  |  | 1.50 |
| CD45hi |  |  | NA | NA | NA | NA | 0.23 | 0.17 | NA | NA | NA | NA | 1.00 |
| CD45+ |  |  |  | 1.39 | 0.71 |  | 0.30 | 0.29 |  | 1.76 |  |  | 0.75 |
| Bcells |  |  |  |  |  |  |  | 0.27 | 0.47 |  |  | 0.47 | 0.50 |
| NK |  |  | 0.67 |  | 0.64 |  | 0.31 | 0.22 | 0.59 | 0.76 |  | 0.66 | 0.25 |
| NKT |  |  |  |  |  |  | 0.19 | 0.31 |  |  |  |  | 0.00 |
| CD11b+ |  |  | 1.62 | 1.62 | 0.71 |  | 0.17 | 0.10 |  | 2.55 |  |  |  |
| Neutr | 0.24 |  | 1.73 | 1.67 | 0.71 |  | 0.06 | 0.06 |  | 2.82 |  |  |  |
| Ly6C- mono/mac |  | 0.55 |  | 1.57 |  |  | 0.51 | 0.40 |  |  |  |  |  |
| Ly6C+ mono/mac |  |  |  |  | 0.71 |  | 0.19 | 0.07 |  |  |  |  |  |
| cDC |  |  |  |  | 0.55 |  | 0.40 | 0.14 |  | 0.42 | 0.66 | 0.58 |  |
| TH |  |  |  |  |  |  | 0.27 | 0.24 |  |  |  | 0.54 |  |
| Treg |  |  |  |  |  |  | 0.30 | 0.18 |  |  |  | 0.50 |  |
| CD8 |  |  |  |  |  |  |  | 0.45 | 0.67 |  |  | 0.52 |  |
| TCRgd |  |  |  |  |  | 0.74 | 0.66 | 0.28 | 0.53 |  |  | 0.55 |  |
| Microglia |  | 0.57 |  |  |  |  |  | 0.54 |  |  |  |  |  |
| Mono |  |  |  |  |  |  | 0.16 | 0.11 |  |  |  |  |  |
| Ly6C+Mono |  |  |  |  |  |  | 0.09 | 0.09 |  |  |  |  |  |
| Ly6C-Mono |  |  |  |  |  |  | 0.54 | 0.16 |  |  |  |  |  |
| Mac |  |  |  |  |  |  | 0.23 | 0.12 |  |  |  |  |  |
| Ly6C+Mac |  |  |  |  |  |  | 0.22 | 0.07 |  |  |  |  |  |
| Ly6C-Mac |  |  |  |  |  |  | 0.45 | 0.64 |  |  |  |  |  |

Two major impacts of the KD on immune cells in the periphery were observed. In the blood at Peak disease, we found that the KD *increased* circulating levels of CD11b**^+^** myeloid cells, most notably neutrophils, in both sexes (Fig. 3A left graph and Table 1). This elevation was sustained at Late disease for males whereas several cell types were decreased in female blood at Late disease, including B cells, NK cells, and some T cell subsets (Fig. 3B and Table 1). Despite these increases in circulating myeloid cells in both females and males at Peak disease, the KD *reduced* or did not change the abundances of these same myeloid populations in the BM at this time point (Fig. 4A and Table 1), indicating that the increased circulating cells were not a result of increased hematopoiesis in the BM.

Overall, these analyses revealed that shared responses among females and males to the KD predominated over differential responses. Paramount among these shared responses were the restoration of motor function with similar kinetics (Fig. 1), the wholesale reduction of infiltrating immune cells in the SC at Late disease preceded by an increase in circulating myeloid cells at Peak disease (Figs. 2 and 3), and numerous other immunological responses to MOG peptide immunization, consistent with published work (5, 7). Because the increased myeloid cells in the blood were not attributable to increased hematopoiesis in the BM (Fig. 4), we hypothesized that the KD promotes blood-spinal cord barrier (BSCB) resealing. This model could at least partially account for the elevated myeloid cells in the blood at Peak disease (Fig. 3A) followed by the dramatic reduction of immune cell infiltrates in the SCs at Late disease (Figs. 2A-D).

### The KD accelerates resealing of the BSCB in symptomatic EAE mice

In both EAE and MS, loss of BSCB integrity is required for select immune cell populations to transmigrate from the circulation into the CNS to promote inflammation and disease onset/progression (reviewed in (21)). Notably, BSCB permeability in the EAE model occurs coincident with symptom onset (22–24). We tested the impact of the KD on BSCB integrity and permeability by injecting male and female EAE mice intraperitoneally with sodium fluorescein (NaFl, molecular weight = 376.3 g/mol) at either 0 DPSO, 3 DPSO, or 6 DPSO. 0 DPSO represents the time at which motor deficits are first observed (i.e., motor score ≥ 1) and mice have not yet been switched from standard chow to the KD or CD. 3 and 6 DPSO represent 3 and 6 days, respectively, of treatment with diet following symptom onset. 30 minutes after NaFl injection, SCs were collected to quantify extravasated fluorescent tracer. The amount of NaFl in the serum of each mouse was also measured to normalize for differences in the volume of tracer injected and/or animal size. These experiments showed that healthy males and females have minimal and comparable BSCB permeability (Figs. 5A and 5B). On the day of motor symptom onset (i.e., 0 DPSO; motor score ≥ 1), the permeability of the BSCB in EAE females was ∼2 fold greater than in males (orange bars in Figs. 5A and 5B, respectively). BSCB permeability peaked at 0 DPSO and was sustained at this level for 3 days in both sexes on CD. Remarkably, by 3 DPSO, the KD decreased the permeability of the BSCB in both females and males by ≥ 50% compared to EAE mice fed the CD (Figs. 5A and 5B). At 6 DPSO, KD-fed males showed a further decrease in BSCB permeability that was comparable to the BSCB permeability found in healthy mice (Fig. 5B). Females at 6 DPSO on both diets had similar permeability, which was greater than that of healthy female mice (Fig. 5A). Notably, both sexes fed the CD for 6 days showed decreased BSCB permeability compared to 0 DPSO due to the spontaneous resealing that occurs with this model (Figs. 5A-B) (22).

As adherens proteins and tight junction proteins maintain BSCB integrity, we performed Western blotting for the adherens junction protein VE-cadherin and the tight junction protein ZO-1 to determine if the KD impacted the levels of either protein at the Peak time point. This analysis showed that the KD increased both VE-cadherin and ZO-1 in females but not in males (Figure 5C-D), consistent with the female-specific reductions of neutrophils (Fig. 2E), IL-6, and CXCL2 (Fig. 2G) that we observed during early symptomatic disease.

Collectively, our interrogation of the BSCB revealed that: (i) there is a diet-independent, demonstrable and spontaneous BSCB resealing in both sexes by 6 DPSO as the symptomatic phase of EAE progresses, (ii) the KD accelerates this BSCB resealing in both sexes but to different extents despite improving motor scores comparably in the mice used for these permeability studies (Supplemental Fig. 3) and (iii) the extent of BSCB permeability at motor symptom onset (i.e., 0 DPSO) is sexually divergent (females > males) as are the KD-mediated increases of VE-cadherin and ZO-1.

A potential mechanism underlying the capacity of the KD to accelerate BSCB resealing could derive from reducing SC levels of secreted (i.e., soluble) IL-1β. This cytokine and its receptor, IL1R1, are required for the progression of EAE by contributing essential roles in BSCB disruption, endothelial cell-neutrophil adhesion, and myeloid cell infiltration into the CNS (22, 25–27). To test this hypothesis, we generated SC extracts from female and male EAE mice and quantified soluble IL-1β by ELISA at the time points matching the NaFl BSCB permeability assay. In females, the KD reduced IL-1β levels in the SC at both 3 and 6 DPSO compared to the CD (Fig. 5E). In males, IL-1β levels at 6 DPSO were reduced in KD versus CD fed mice and at 3 DPSO, trended downward for KD-fed mice but did not reach statistical significance although the levels in the KD group (but not the CD group) at this time point were statistically significantly lower compared to 0 DPSO (Fig. 5F). As expected, IL-1β levels at all time points for both sexes and diets were markedly elevated in EAE mice compared to healthy mice (Figs. 5E-F), and the kinetics of BSCB permeability and resealing (Figs. 5A-B) mirrored the directional change in the levels of soluble IL-1β (Figs. 5E-F).

To determine if the KD retained efficacy and promoted BSCB resealing when initiated after symptoms had persisted, we repeated the NaFl tracer studies but waited to initiate feeding of the therapeutic diet until 6-8 DPSO (after all animals reached a motor score of ≥ 2.5). The fluorescent tracer was injected 14-16 days after initiating the KD or CD (i.e., 20-23 DPSO) (Fig. 6A). Remarkably, the KD retained the capacity to improve motor scores when initiated ∼ 1 week after clinical symptom onset, with functional improvements detected within 6 days of treatment for females (Fig. 6B) and 4 days for males (Fig. 6D). Additionally, in both sexes, the KD resealed the BSCB compared to the permeability on the day of diet switch (Figs. 6C and 6E). In males, this resealing was also statistically significant when compared to CD-fed mice (Fig. 6E).

We hypothesized that the reduction of soluble IL-1β in the SCs by the KD was a consequence of SC infiltrates either: (i) synthesizing less precursor IL-1β, (ii) producing less secreted IL-1β via inhibition of caspase-1/inflammasome-mediated proteolytic processing of the precursor cytokine, and/or (iii) undergoing increased apoptosis. We therefore simultaneously measured total intracellular levels of IL-1β (precursor + soluble but not-yet-secreted), caspase 1-inflammasome activity, and apoptosis in SC infiltrating and resident immune cells recovered from EAE mice fed the CD or KD for either 2-4 days or at Peak disease. Notably, prior work had demonstrated that secreted IL-1β is primarily produced by the proteolytic activity of inflammasome-associated caspase-1, and that caspase-1 is required for the loss of blood-CNS barrier integrity in the EAE model (26, 27). Additionally, the ketone body, β-HB, has been shown to suppress caspase-1/inflammasome activity (e.g., (28, 29)) and either induce (e.g., (30)) or inhibit (e.g., (31, 32)) apoptosis depending on the cell type and context. Isolated SC immune cells were subjected to intracellular IL-1β labeling with an antibody that recognizes precursor and mature (soluble) cytokine, fluorochrome inhibitor of caspases (FLICA) labeling to assess caspase-1/inflammasome activity, and with the apoptosis marker, Annexin V, as well as with a cell viability dye to mark dead cells. FLICA is a flow cytometry assay that uses a cell permeable, fluorescently-labeled inhibitor peptide to selectively bind activated caspase-1 in live cells as a readout of inflammasome activity (33).

Unexpectedly, at 2-4 DPSO the KD did not significantly change the percentage of FLICA+ and IL-1β + cells in the SC (data not shown) but at Peak disease, the diet *increased* both markers across multiple cell types (Supplemental Fig. 4A-B). Yet, the KD increased the percentage of numerous cell types that were positive for Annexin V labeling, consistent with amplified cell death via apoptosis at both time points in both sexes (Fig. 7). For females, this increased apoptosis was observed at 2-4 DPSO for neutrophils, Ly6C**^+^** mono/macs, Ly6C**^-^**mono/macs, NK cells, and NKT cells (Fig. 7A). For males at 2-4 DPSO, increased Annexin V was observed in Ly6C**^+^** mono/macs, Ly6C^-^ mono/macs, and TH cells (Fig. 7A). At Peak disease, both females and males shared increases in Annexin V positive cells for microglia, Ly6C**^+^**mono/macs, Ly6C^-^ mono/macs, NKTs, TH cells, CD8+ T cells, and TCR-γδ T cells (Fig. 7B). These data suggest that the accelerated BSCB resealing promoted by the KD decreases soluble IL-1β as well as the abundance of infiltrating cells in the SC by increasing apoptosis of multiple SC infiltrating cell types and of resident microglia rather than by inhibiting the inflammasome in these cells.

**Figure 7.**
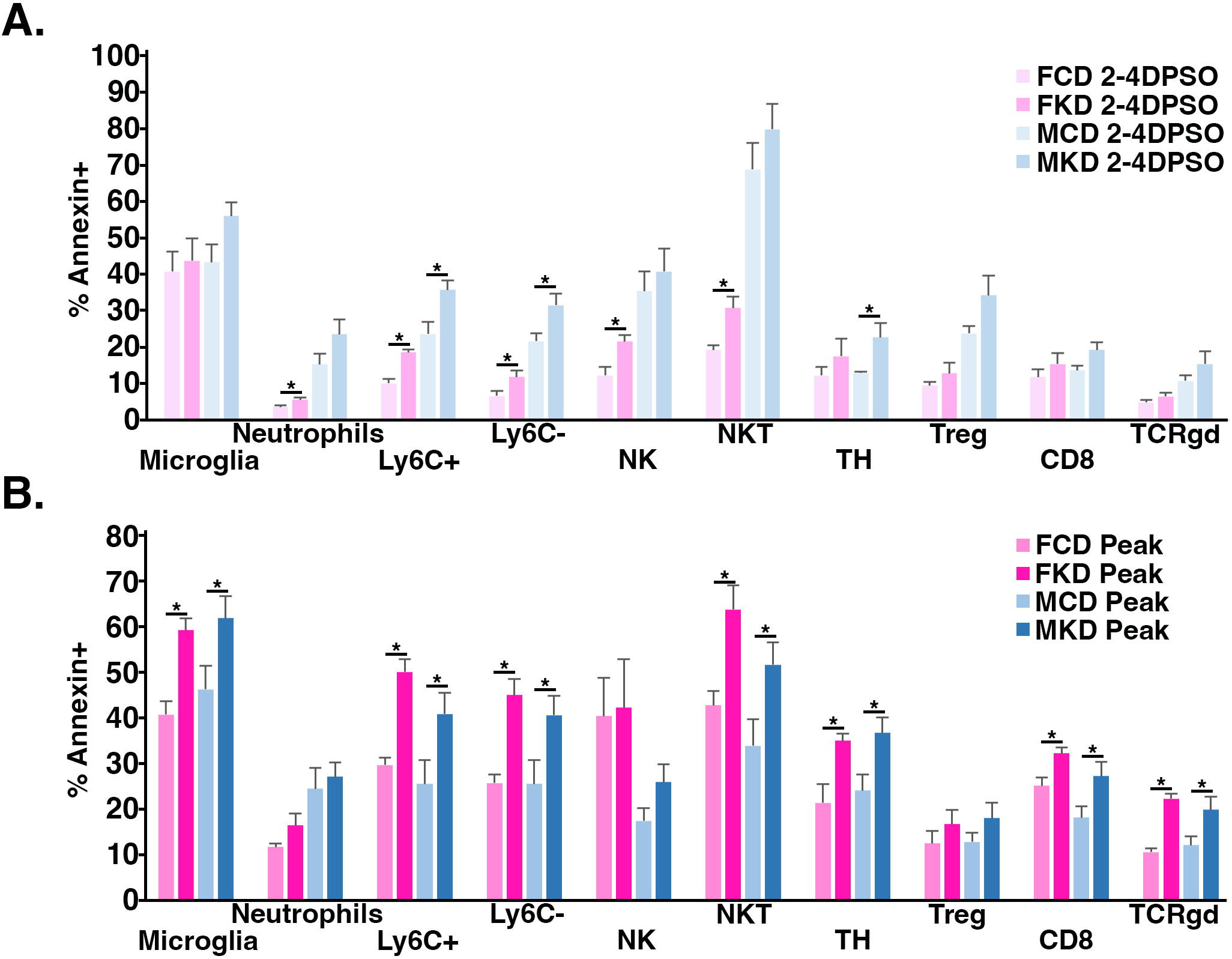
KD increases the percentage of apoptotic immune cells in the CNS. Flow cytometry analysis of SC at (**A**) 2-4 DPSO and (**B**) Peak (5-7 DPSO) to determine the percent of specific cell types positive for Annexin V as a marker of apoptosis. n=6-10 mice/sex/diet/timepoint. Asterisks in all graphs denote statistical significance *p<0.05 vs CD as determined by two-tailed, unpaired Student’s t test.

As neutrophils are primary drivers of blood-CNS barrier loss (34), are localized at sites of blood-brain barrier (BBB) leakage in MS and *neuromyelitis optica spectrum disorder* postmortem samples (e.g., (22)) and were most impacted by the KD (Table 1), we further characterized them at both Peak and Late Disease. t-SNE analysis showed a striking KD-dependent dropout of neutrophil abundance at Late disease in EAE female and male SCs with a notable retention of select neutrophil subpopulations (see black arrows in Fig. 2B and Figs. 8A-B). Notably, neutrophils are defined by being CD11b**^+^**, Ly6C**^int^**, and Ly6G**^+^**as shown in the heatmap overlays (Figs. 8A-B lower panels). To distinguish the retained subpopulations from the bulk of neutrophils that were lost, and to test the hypothesis that the persistent neutrophil subpopulations have an anti-inflammatory (or naïve) phenotype, we analyzed the differential expression of select surface makers. Heatmap overlays of the t-SNE visualization showed that the neutrophils preserved in the SCs of both sexes had reduced expression of Ly6G (Figs. 8A-B lower panels and Fig. 8C). This neutrophil-defining glycoprotein modulates neutrophil migration to sites of inflammation (35), and decreases in the median fluorescence intensity (MFI) of Ly6G have been shown to be a distinguishing marker of pro-resolution neutrophils in an optic nerve crush model (36). Quantification of SC neutrophil Ly6G MFI by flow cytometry confirmed that in both females and males the KD decreased Ly6G expression levels (Fig. 8C). In contrast, Ly6G MFI was unchanged on neutrophils in the blood (Supplemental Fig. 4C) revealing an anatomical compartment-specific effect of the KD in EAE mice. The reduction of Ly6G MFI in neutrophils by the KD was additionally detected at Peak disease indicating that it is initiated early, before consistent and dramatic neutrophil loss was observed in males and then sustained through Late disease in both sexes (Fig. 8C).

**Figure 8.**
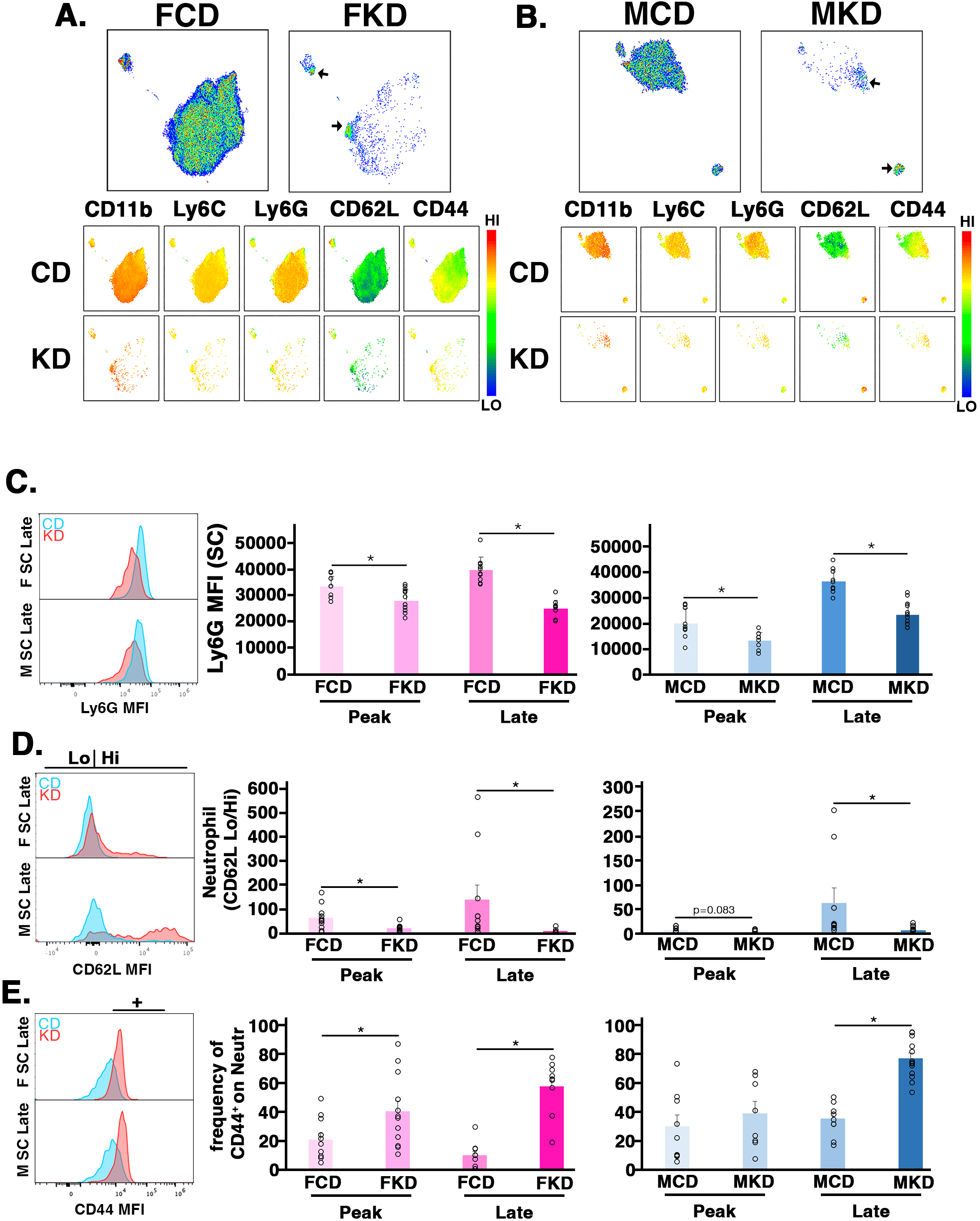
The KD reduces total neutrophil abundance in the SCs of EAE mice but preserves subpopulations expressing surface markers associated with inflammation resolution. (A,B) Top row: t-SNE visualization of total neutrophils in the SCs of female (A) and male (B) EAE mice treated with CD or KD and harvested at Late disease taken from Fig. 2B. Black arrows on FKD and MKD plots mark neutrophil populations retained in the KD-fed mice. The sets of 10 smaller t-SNE plots show intensity of labeling of neutrophils for the indicated cell surface markers for females (left set) and males (right set). Heat map indicates HI (red) and LO (blue) expression. (**C**) Representative histogram and graphs of Ly6G median fluorescence intensity (MFI) on SC neutrophils as a function of sex, diet, and time of disease course. (**D**) Representative histogram of CD62L MFI and graphs of SC neutrophil activation states represented by the ratio of CD62L^LO^ to CD62L^HI^ expression determined by flow cytometry. (**E**) Representative histogram of CD44 MFI and graphs of frequency of CD44**^+^** on SC neutrophils. Open circles represent individual animals. n=9-13 mice/sex/diet. Asterisks in all graphs denote statistical significance *p<0.05 as determined by two-tailed, unpaired Student’s t test or alternatively, by Mann-Whitney U test if Shapiro-Wilk test for normality showed a non-normal distribution. Data for females shown in left graphs and for males in right graphs.

The t-SNE plots also revealed that the small, retained cluster of neutrophils separating from the main cluster were enriched for CD62L and CD44 (Figs. 8A-B lower panels and Figs. 8D-E). CD62L is an adhesion molecule also known as ‘L-selectin’ that plays critical roles in the homing and chemotaxis of lymphocytes to lymphoid organs as well as to sites of inflammation and injury (reviewed in (37)). Neutrophils typically shed CD62L upon being activated and as an outcome of transmigration into inflamed tissues (reviewed in (38)), though loss of the marker is not strictly required for such infiltrating activity (e.g., (39)). Thus, the presence of CD62L is associated with naïve homeostatic neutrophils whereas loss of the marker is typically linked to neutrophil activation during inflammation. Using the ratio of CD62L**^LO^** to CD62L**^HI^** as a proxy for neutrophil activation state with a higher ratio indicating greater activation (e.g., (40)), we found that the KD dramatically reduced the ratio of CD62L**^LO^** to CD62L**^HI^**in the SCs of both sexes, consistent with the retained neutrophils having reduced pathogenicity (Fig. 8D).

Akin to CD62L, the homing receptor CD44 functions in the adhesion of neutrophils to the endothelium and to their subsequent migration out of blood vessels to enter inflamed tissue (41, 42). CD44 can be internalized from the membrane of activated neutrophils and enter the nucleus to modulate azurophilic granules (43). The t-SNE plots showed that the neutrophils remaining in the SC after KD treatment were enriched for surface expression of CD44 in both sexes (Figs. 8A and B, lower panels). We verified this by showing that the KD increased CD44 MFI in the SC of both sexes as well as the frequency of neutrophils expressing CD44 on their surface at Peak disease in females and at Late disease in both females and males (Fig. 8E). Together, these data indicate that the neutrophils remaining in the SC at Late disease for mice fed the KD are phenotypically distinct (i.e., Ly6G**^LO^**, CD62L**^HI^**, CD44**^+^**) from the pro-inflammatory neutrophils (i.e., Ly6G**^HI^**, CD62L**^LO^**, CD44**^-^**) that drive motor deficits in EAE.

A second striking impact of the KD in the SC was that it severely reduced the abundance of mono/macs expressing Ly6C from the progression of Peak to Late disease in both sexes (i.e., > 70% in females and > 90% in males) whereas for CD-fed mice, the abundances did not change (Supplemental Fig. 5A). In contrast to the loss of Ly6C**^+^** mono/macs, the abundances of Ly6C**^-^** mono/macs from Peak to Late did not significantly change as a function of either diet (Supplemental Fig. 5B). Expression of the Ly6C glycoprotein by mono/macs plays an integral role in EAE with higher expression on infiltrating cells associated with pro-inflammatory phenotypes and lower expression associated with tissue repair (44). The KD-driven reduction of Ly6C across mono/macs was likewise illustrated by t-SNE analyses comparing mono/macs between diets at Late disease and normalized for cell number (Fig 9A). This reduction was also manifested when we analyzed Ly6C MFI (Fig. 9B histograms) and by graphing the frequency of mono/macs that are Ly6C**^+^** as a function of time and diet in both sexes (Fig. 9B graphs). These data indicate that the KD skews myeloid cell populations (e.g., neutrophils, mono/macs) in the CNS of EAE mice away from pathogenicity and in favor of resolving inflammation and promoting tissue repair/recovery.

**Figure 9.**
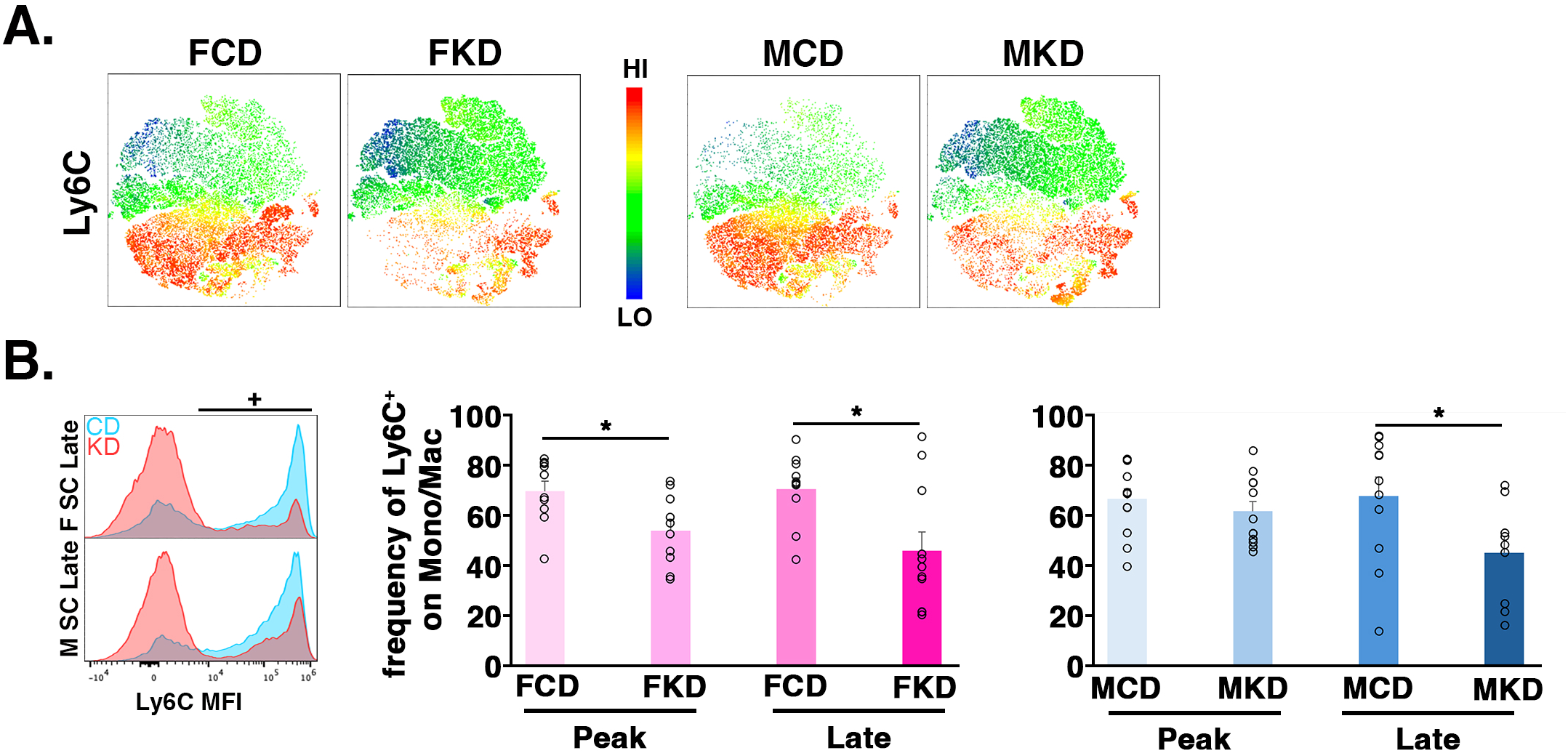
The KD reduces mono/macs with high Ly6C expression in the SCs of EAE mice but preserves subpopulations associated with resolving inflammation. **(A)** t-SNE visualization comparing equal numbers of mono/macs in the SCs of female and male EAE mice treated with CD or KD and harvested at Late disease. Heatmap overlay shows KD-mediated reduction in cells with high Ly6C expression. **(B)** Representative histogram of Ly6C MFI on mono/macs at Late disease along with a graph showing KD-mediated reductions in the frequency of mono/macs that are Ly6C**^+^** in the SC at Peak and Late disease for both sexes. Open circles represent individual animals. n=9-13 mice/sex/diet. Asterisks in all graphs denote statistical significance *p<0.05 vs CD as determined by two-tailed, unpaired Student’s t test.

We next tested the efficacy of the KD in a global nicotinic acid receptor (a.k.a., hydroxycarboxylic acid receptor 2 (HCA**_2_**) or GPR109A) knockout strain undergoing EAE. HCA**_2_** is expressed on neutrophils, dendritic cells, macrophages, Langerhans cells, retinal pigment epithelial cells, keratinocytes, white and brown adipocytes and microglia with more limited expression on activated lymphocytes (45, 46). Our rationale was twofold. First, β-HB is an agonist of the HCA**_2_**, and multiple lines of evidence show that activation of the receptor by β-HB inhibits NF-ΚB signaling upstream of the NLRP3 inflammasome (47–49) and restricts neutrophils from accumulating in the CNS (50). Interestingly, the receptor is required on neutrophils for the efficacy of the MS drug, dimethyl fumarate (DMF) (50, 51). Second, akin to the KD (Fig. 9), HCA**_2_**-mediated signaling has been associated with reduced Ly6C expression on mono/macs (reviewed in (52)). Unexpectedly, the efficacy of the KD to restore motor function and reduce SC infiltrates was not diminished in mice genetically ablated for HCA**_2_** (Supplemental Figs. 6A-C).

Lastly, we tested the requirement of nuclear factor erythroid 2-related factor 2 (Nrf2) for KD-mediated efficacy in the EAE model. Nrf2 is a transcription factor that induces the expression of target genes in response to oxidative, nitrosative, and multiple other types of stress (reviewed in (53, 54)). The protein products of Nrf2 target genes in turn restore and maintain redox and proteome homeostasis. Notably, the benefits of KDs in some rodent disease models of neurodegeneration have been shown to involve Nrf2 activation and the accompanying expression of Nrf2 target genes (e.g., Alzheimer’s mice (55, 56)), SC injury in rats (57, 58); toxin-induced Parkinson’s mouse model (59)). Akin to the HCA_2_ knockout data, genetic ablation of Nrf2 did not detectably impact the efficacy of the KD to restore motor function (Supplemental Fig 6D-E). Notably, when comparing the impact of the CD or the KD in either knockout strain versus its strain-matched wild type counterpart, average daily motor scores were comparable (Supplemental Figs. 6B and 6E, respectively). Together, these knockout studies show that HCA_2_ and Nrf2 are dispensable for the functional recovery and efficacy of our MCT, ω3-fatty acid, and fiber-enriched KD in the interventional EAE paradigm.

## DISCUSSION

Small clinical trials involving KDs and low glycemic diets (e.g., the Wahls diet) for pwMS have shown promising outcomes including reduced fatigue, improved quality of life, improved sleep quality, reduced expanded disability status scale scores, reduced levels of serum neurofilament light chain and markers of inflammation, and improvements in 6-minute walk test and nine-hole peg tests (60–66). However, the mechanisms underlying these therapeutic outcomes remain largely unknown. Toward the goal of identifying such mechanisms, and in agreement with recent studies in rodents showing that a KD (by raising β-HB levels) reduces T cell pathogenicity (12, 67–69), we found that the KD reduced T cells in the SC at Late disease by >70% (Table 1). Two major features distinguish our work from and complement these prior studies. First, we started feeding the KD at symptom onset whereas the previous studies fed the diet either 10 days prior to (67) or coincident with (69) MOG immunization. Secondly, our mechanistic analyses primarily interrogated the impact of the KD on myeloid cells in both sexes whereas the previous studies focused on lymphocytes with most experiments being done in only a single sex.

Our study advances the field in several ways. We discovered that the KD accelerated the kinetics of BSCB resealing (Figs. 5 and 6). Underlying this effect was reduced SC levels of soluble IL-1β the primary cytokine driving loss of BSCB integrity (27, 70). Additionally, this accelerated BSCB resealing and improvement of motor scores occurred whether the KD was started the day of symptom onset (Fig. 5 and (11)) or a week after symptom onset (Fig. 6). For both treatment regimens and in both sexes, the KD began restoring motor functions within a few days, indicating the potential to leverage this dietary strategy to mitigate symptoms in response to acute relapses for pwMS and for treatment-naïve individuals experiencing clinically isolated syndrome prior to a definitive MS diagnosis. Third, in both sexes, the KD robustly decreased the abundance of total SC infiltrates, at least in part by increasing the proportion of apoptotic cells (Fig. 7), while preserving subpopulations of neutrophils and mono/macs expressing markers consistent with anti-inflammatory, pro-resolution phenotypes (Figs. 8 and 9, respectively). Lastly, we found that the efficacy of the KD when used as an interventional treatment was independent of HCA_2_ and Nrf2 (Supplemental Fig. 6), two proteins at the center of signaling nodes linked to the beneficial effects of β-HB (e.g.,(71–73)).

In EAE, both neutrophil and mono/mac transmigration into the CNS parenchyma is accompanied by an amplification of inflammasome activity that results in IL-1β secretion, which in turn drives BSCB integrity loss (22, 26). Thus, the reduction of soluble IL-1β levels in the SC by the KD (Figs. 5E-F) supports that the accelerated restoration of BSCB integrity could stem from β-HB-mediated inhibition of the NLRP3 inflammasome (29) or inhibition of signaling upstream of NLRP3 inflammasome activation (e.g., P2X7 receptor activity (74, 75)). Additionally, the KD could promote apoptosis of infiltrated myeloid cells to limit these primary sources of IL-1β production in the CNS. Studies to determine if either, or both, of these potential mechanisms were contributing to the reduction of IL-1β revealed that the KD increased the fraction of multiple infiltrating cell types undergoing apoptosis (Fig. 7) but did not inhibit caspase-1/inflammasome activity (Supplemental Figs. 4A-B). The data from our IL-1β, FLICA, and Annexin V analyses together are consistent with the KD reducing soluble IL-1β in the SC by promoting or sustaining apoptosis of infiltrating cells while simultaneously reducing the influx of new infiltrates through limiting BSCB permeability. It is also possible that the KD directly promotes apoptosis of one or more pathogenic cell types by increasing the expression of pro-apoptotic mediators and/or reducing the expression of anti-apoptotic factors (e.g., (30–32)).

Additional mechanistic contributions could come from β-HB promoting BSCB resealing by (i) increasing the expression of tight junction and adherens junction proteins among the endothelial cells that comprise the barrier (Fig. 5C-D), and/or (ii) reducing the expression of matrix metalloproteinases (MMPs), like MMP-9, that degrade CNS-blood barriers (e.g., (76, 77)). These activities stem from β-HB inhibiting histone deacetylase 3 (HDAC3) (78, 79) coupled to HDAC inhibition decreasing inflammasome activation (80), and are consistent with what was reported for an MCT-enriched KD (81).

Neutrophils were the earliest and most sustainably impacted cell type in response to the KD (Figs. 2 and 8). These granulocytes have been relatively underappreciated in MS due to their limited detection in mature lesions, but compelling evidence implicates them in the acute stages of lesion formation in both MS and EAE (22). Further, neutrophils are elevated and primed in the blood and cerebral spinal fluid during MS relapses (82, 83), and administration of the neutrophil recruitment factor G-CSF severely exacerbated disease in some pwMS (84). The KD significantly reduced neutrophil abundance in female SCs after only a few days of diet treatment (Fig. 2E) and dramatically decreased neutrophils (i.e., by > 90%) in the SCs of both sexes at Late disease (Fig. 2C), likely through accelerated BSCB resealing. These reductions fit with the central role that neutrophils, in conjunction with microglia, astrocytes, and endothelial cells, play in mediating blood-CNS barrier breakdown in EAE (22, 85–87) and other models of CNS injury (87, 88). Moreover, the robust depletion of neutrophils by the KD in females at Peak disease coincided with reductions in SC levels of IL-6 and the neutrophil chemoattractant CXCL2, which function cooperatively in recruiting neutrophils to sites of inflammation (Fig. 2G). Remarkably, the neutrophils that persisted in the CNS of KD-fed mice had surface marker expression profiles favoring anti-inflammatory and pro-resolution phenotypes; these were evident at both Peak and Late for females but with the exception of Ly6G MFI, only at Late disease for males (Fig. 8).

In addition or related to cell death by apoptosis (Fig. 7), the dropout of myeloid cell populations expressing pro-inflammatory markers in the SCs of KD-treated mice (Figs. 8 and 9) may involve the reliance of these cell types on metabolic reprogramming that favors glycolysis for their activation, expansion, and eventual pathogenicity (reviewed in (44, 89)). For neutrophils, restricting glycolysis within the hypoxic environment of inflamed tissues can increase apoptosis (90), and this effect is cell autonomous as confirmed by culturing neutrophils in limited glucose (91). The severe dietary carbohydrate restriction imposed by the KD may therefore amplify neutrophil apoptosis at sites of inflammation within the CNS.

The KD-mediated reduction of classical inflammatory (Ly6C^+^) (Fig. 9) mono/macs was accompanied by a preservation of non-classical pro-resolution (Ly6C^-^) mono/macs (Supplemental Fig. 5B). Notably, a bioenergetic shift underlies the phenotypic switching of monocytes from Ly6C**^+^**to Ly6C**^-^** that fits with the very low carbohydrate content of the KD and could in part explain these findings. Specifically, Ly6C**^+^** monocytes rely on enhanced glycolysis mediated by IL-17 and GM-CSF driven upregulation of glucose transporters to import more glucose (92) whereas a switch of Ly6C**^+^** monocytes that favors oxidative phosphorylation promotes conversion to non-classical Ly6C**^-^**mono/macs involved in inflammation resolution (93). Similar reductions of Ly6C**^+^** mono/macs and phenotypic switching of the remaining mono/macs from pro-inflammatory to anti-inflammatory has been observed for pharmacological and genetic strategies that mitigate EAE and other murine models of CNS demyelination (e.g., (94)). For example, reductions of Ly6C**^+^** monocytes leading to ameliorated EAE have been reported for nucleotide-oligomerization domain 2 (NOD2) agonists (94), C-C chemokine receptor 4 (CCR4) knockout mice (95), and IL-17 knockout mice (96). Future studies to determine if the KD shares mechanisms of efficacy with the above could provide additional insights into this dietary intervention. Collectively, these findings together with those in the literature support that acute implementation of dietary carbohydrate restriction could provide a relatively safe, rapid, and reversible means of tempering myeloid cell pathogenicity during autoimmune flare-ups.

Our flow cytometry analyses of SC, blood, and BM showed that the KD differentially impacted the immune cell profiles of female and male EAE mice to a modest degree in the SC, but more extensively in the blood and BM (Figs. 3 and 4, respectively). Many of these peripheral immune cell changes were also observed in healthy age-matched mice fed the KD for 2 weeks. Thus, these changes are not EAE disease-specific (Supplemental Table 4) and therefore were not a focus of the current study. Although the implications of these sex differences in peripheral immune cells remain to be determined, their collective effects appear to be relatively minor compared to the major *shared* impacts of the KD to: (i) restore motor deficits comparably in the sexes (Fig. 1), (ii) promote BSCB resealing in both sexes (Figs. 5 and 6), (iii) severely reduce the abundance of myeloid and T cell infiltrates in the CNS of both females and males (Table 1), (iv) increase the percentage of apoptotic cells in the SC (Fig. 7), and (v) preserve neutrophil and mono/mac subpopulations associated with resolving inflammation in the SC of both sexes (Figs. 8 and 9).

In conclusion, this work advances novel mechanisms by which a KD mitigates autoimmune-mediated MS-like pathologies to promote functional recovery (Fig. 10). These findings offer a plausible and unifying explanation for the beneficial impacts of KDs across a range of CNS diseases involving disrupted blood-CNS barriers and myeloid cell-mediated pathogenicity including MS, drug-resistant epilepsy, Amyotrophic lateral sclerosis, Alzheimer’s disease, Parkinson’s disease, and schizophrenia (e.g., (61, 65, 97–101)).

**Figure 10.**
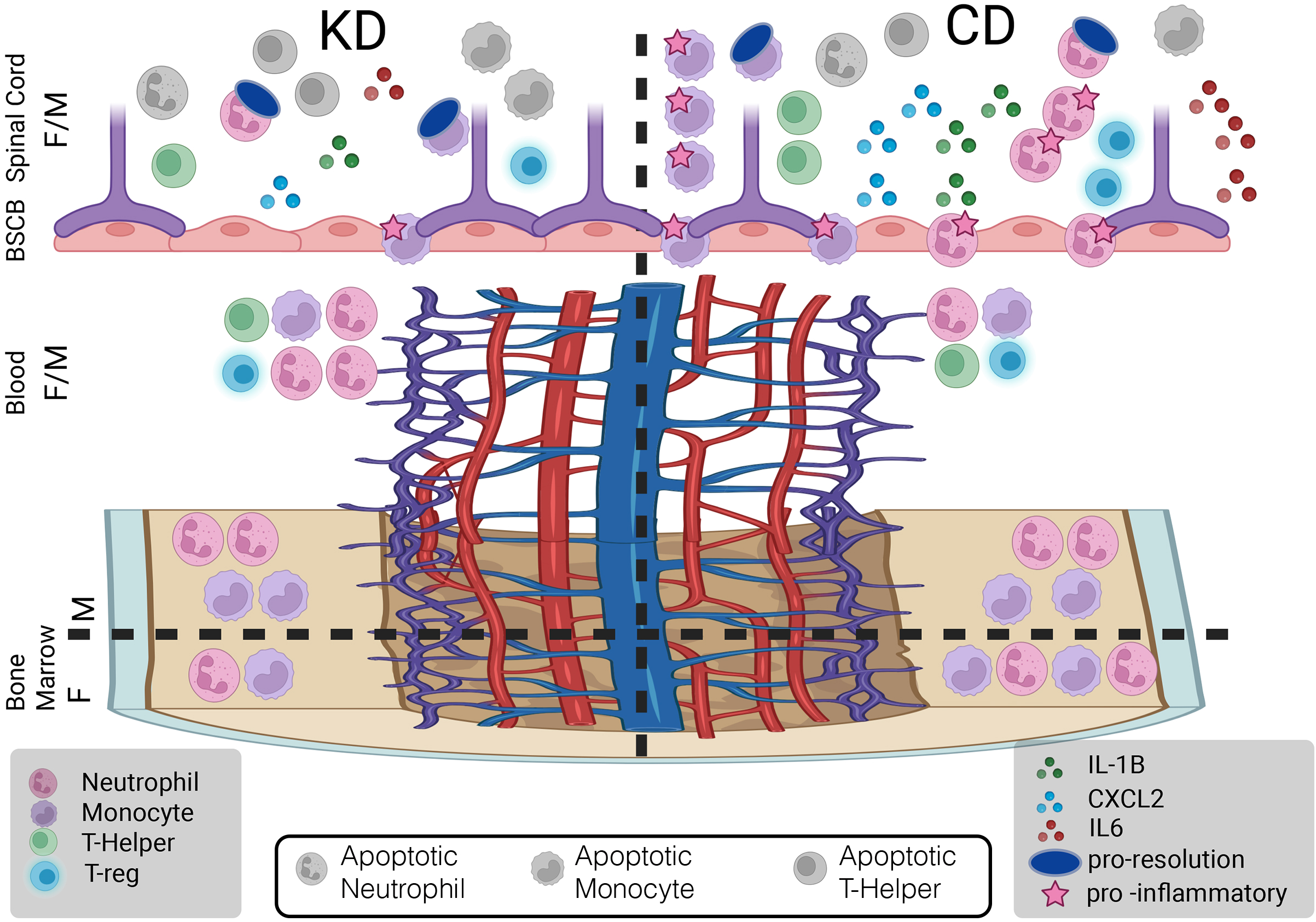
Model of the impacts of the KD on immune cells and the BSCB. Our data support a model in which the KD restores motor function to symptomatic EAE mice by (i) accelerating resealing of the BSCB to limit the accumulation of neutrophils and monocytes in the SC, (ii) limiting T cell accumulation in the CNS and increasing apoptosis of multiple cell types, (iii) reducing cytokines in the SC that drive the loss of BSCB integrity (IL-1β) and recruit neutrophils to sites inflammation (IL-6 and CXCL2), (iv) causing an early accumulation of myeloid cells in the blood and (v) preserving myeloid cells that promote inflammation resolution while limiting myeloid cells that foster inflammation.

## ACKNOWLEDGEMENTS

This work was supported in part by grant R01EY033782 from the National Eye Institute of the National Institutes of Health. The content is solely the responsibility of the authors and does not necessarily represent the official views of the National Institutes of Health. We thank Tamy Aguero-Fraire for technical assistance, and Dr. Katarzyna Zyla-Jackson for help with harvesting female SCs and staining female SC cells for flow cytometry during initial studies. We thank Jacob Bass in the OMRF Flow Cytometry Core and Dr. Caleb Marlin for the Axioscan analysis using the instrument purchased by the ACI program at OMRF. We are grateful to Dr. Stefan Offermanns (Max Planck Institute for Heart and Lung Research, Germany) for permission to obtain the HCA**_2_** knockout mice that his laboratory created.

## AUTHOR CONTRIBUTIONS

KSP: experimental design, acquisition and analysis of data, writing, editing. DAW: acquisition and analysis of data, editing. NP: model design from data analysis. SK and RCA: data analysis and editing. SMP: experimental design, data analysis, writing, and editing.

## DATA AVAILABILITY

The data generated and analyzed for this study are all included in this published article and its Supplementary information files.

---

**Supplemental Figure 1. Th**e **KD reduces lesion size and cellularity in the spinal cords of EAE mice.** Spinal cords were harvested at the Late time point and processed for IHC with anti-PLP (the main protein component of myelin) to mark myelin loss and with Hoechst to mark cellularity as a read-out of cellular infiltrates and resident microglia. Insets show lesion outline (PLP) and extent of cellularity (Hoechst).

**Supplemental Figure 2. Flow cytometry gating strategies.** Gating strategy used on cells for immune cell profiling. SC gating strategy shown on top; blood and BM strategy shown on the bottom.

**Supplemental Figure 3. Th**e **KD improves motor function comparably in both sexes.**Graph of EAE motor scores for mice used in the NaFl studies shown in Fig 5. Sex and diets indicated in the key across the top. Asterisks denote statistical significance *p<0.05 as determined by two-tailed, unpaired Student’s t test.

**Supplemental Figure 4. The KD increases inflammasome activity within multiple cell types but does not change the Ly6G MFI of circulating neutrophils. (A, B)** Graphs of flow cytometry data of SC at Peak (5-7DPSO) showing the percent of each listed cell type positive for (**A**) FLICA and (**B**) IL-1β; both are readouts of inflammasome activity. (**C**) Graphs of Ly6G MFI on blood neutrophils as a function of sex, diet, and time of disease course. Open circles represent individual animals. n=9-13 mice/sex/diet. *p<0.05 as determined by two-tailed, unpaired Student’s t test. ns = not statistically significant.

**Supplemental Figure 5. Th**e **KD reduces Ly6C^+^ mono/mac abundances but preserves Ly6C^-^ mono/macs from Peak to Late disease in the SC.** Graphs of abundances of Ly6C**^+^ (A)** and Ly6C**^-^ (B)** mono/macs isolated from SCs at Peak and Late disease. n=9-13 mice/sex/diet. Asterisks denote statistical significance between Peak and Late disease. *p<0.05 as determined by Mann-Whitney U test. ns = not statistically significant.

**Supplemental Figure 6. HCA2 and Nrf2 are not required for the efficacy of the KD in the interventional paradigm. (A)** Motor scores of CD and KD fed female and male HCA_2_ KO EAE mice in the interventional paradigm. **(B)** Average daily motor score of WT and HCA_2_ KO EAE mice are comparable. **(C)** Fold change in number of indicated infiltrating and resident immune cells. Only significant values as determined by Mann-Whitney U test are shown with intensity of green heatmap indicating degree of suppression. Blank boxes indicate not statistically significant between KD and CD for a given cell type. **(D)** Motor scores of CD and KD fed female and male Nrf2 KO EAE mice in the interventional paradigm. **(E)** Average daily motor scores of WT and Nrf2 KO EAE mice are comparable. Open circles represent individual animals. n=5-9 mice/sex/diet/strain. *p<0.05 as determined by two-tailed, unpaired Student’s t test. ns = not statistically significant.

**Supplemental Table 1**: List of fluorochrome-labeled antibodies, companies, and catalog numbers used for flow cytometry studies.

**Supplemental Table 2.** Name, abbreviation and flow cytometry surface markers used to identify each immune cell type analyzed.

**Supplemental Table 3. Immune cell abundances for SC, blood, and BM of KD-fed and CD-fed EAE mice.** Average (avg) cell counts and standard error of the mean (sem) are provided for EAE mice harvested at Peak and Late from SC, blood, and BM. Data for healthy (non-EAE) age-and sex-matched mice fed the KD or CD for two weeks is also provided. Statistically significant differences (* p<0.05) determined by two-tailed, unpaired Student’s t test for blood and BM or by Mann-Whitney U test for SC are bolded.

**Supplemental Table 4. Immune cell profiling for healthy animals fed the KD for 2 weeks.** Tissues were harvested from blood and BM for immune cell profiling. Table lists fold change of KD compared to CD, and only those cell types that reached statistical significance are assigned fold change. Statistically significant differences determined by two-tailed, unpaired Student’s t test. Blank cells indicate that statistical significance was not reached for that cell type. Heatmaps show **green** entries representing fold decreased cell abundance for KD and **red** entries representing fold increased cell abundance for KD. The raw data to generate this table are provided in Supplemental Table 3.

