## Supplement Figures and Tables for "A KETOGENIC DIET ACCELERATES BLOOD-SPINAL CORD BARRIER RESEALING AND DECREASES SPINAL CORD IMMUNE CELL ACCUMULATION IN A PRECLINICAL MODEL OF MULTIPLE SCLEROSIS"

### Supplemental Figure 1

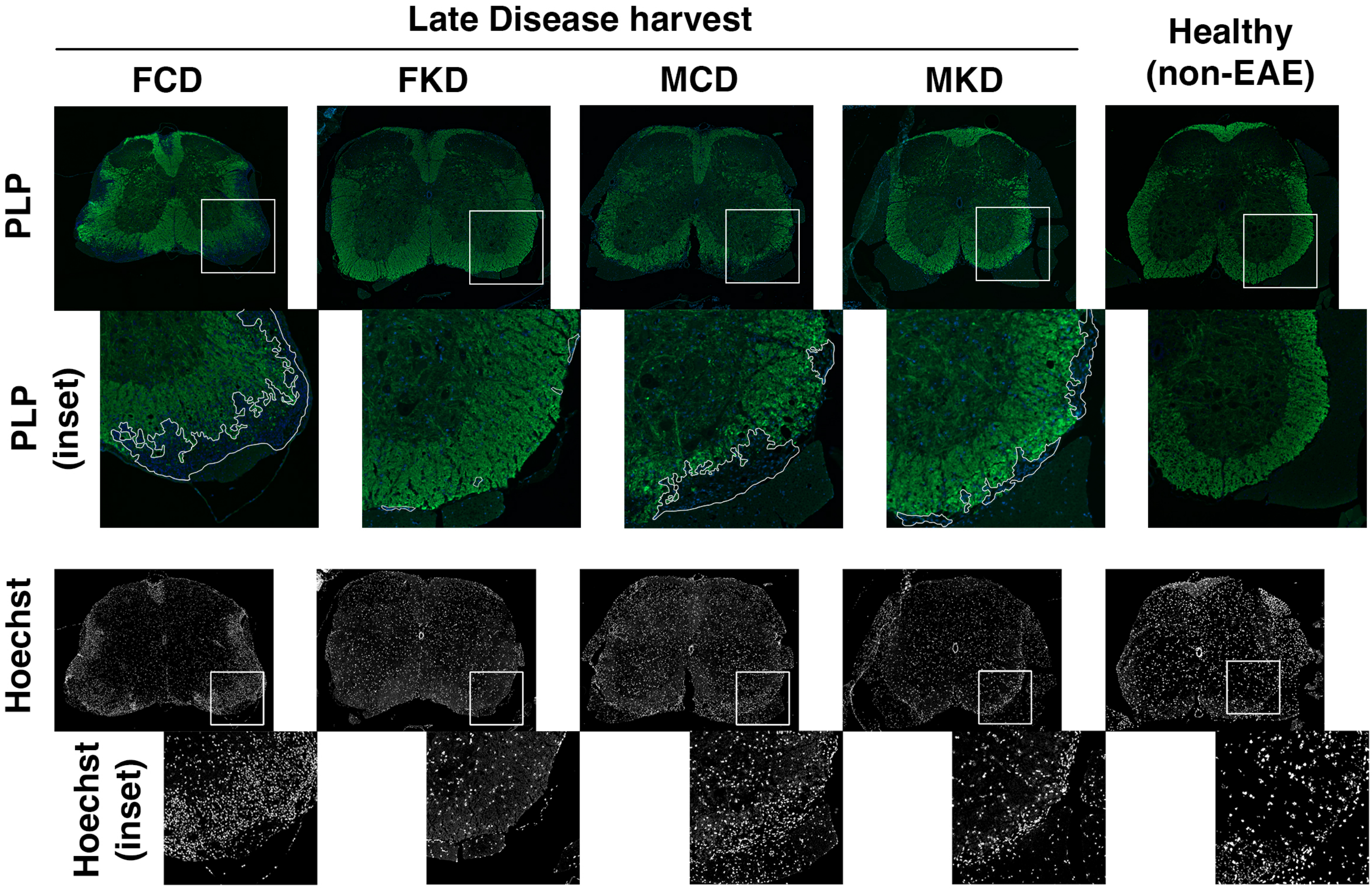

### Supplemental Figure 2

## SC

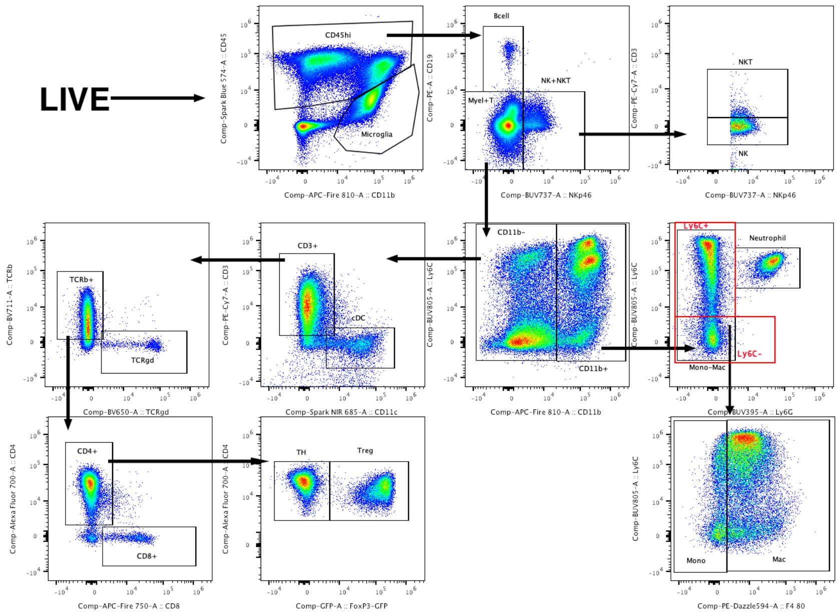


#### BLOOD and BM

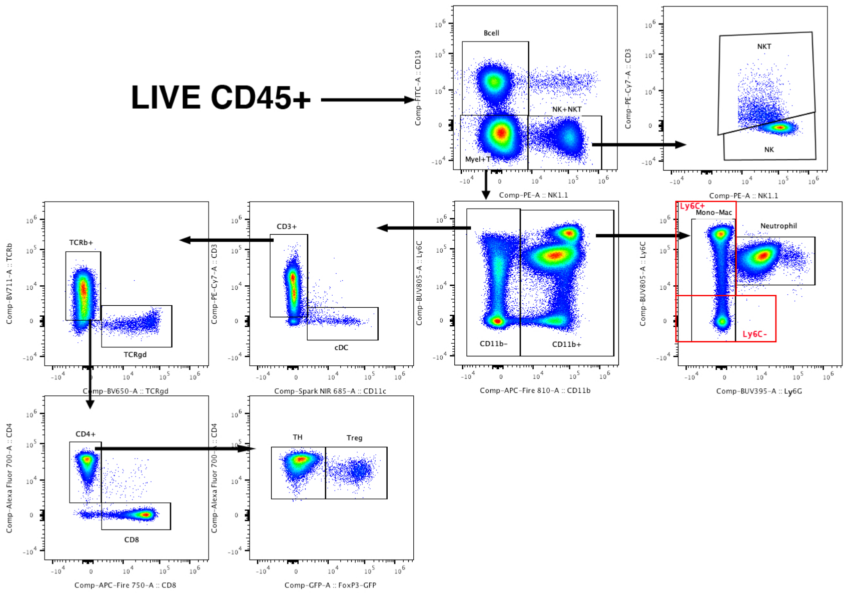

### Supplemental Figure 3

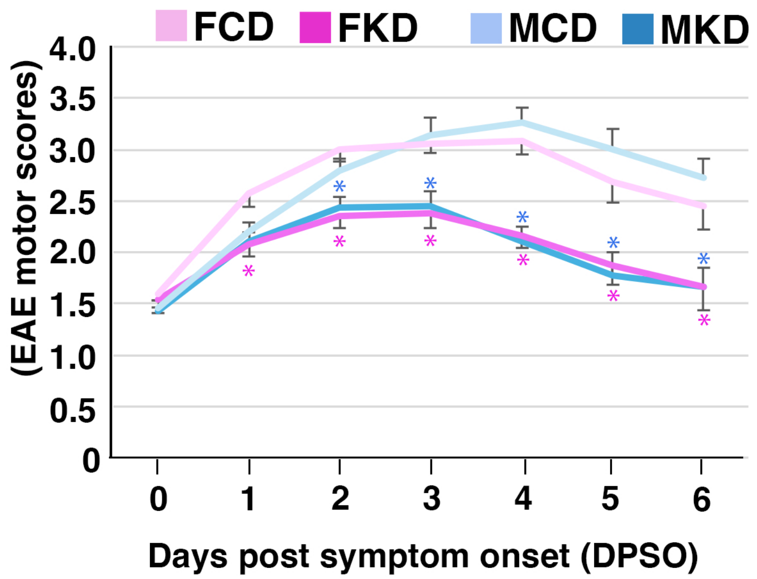

### Supplemental Figure 4

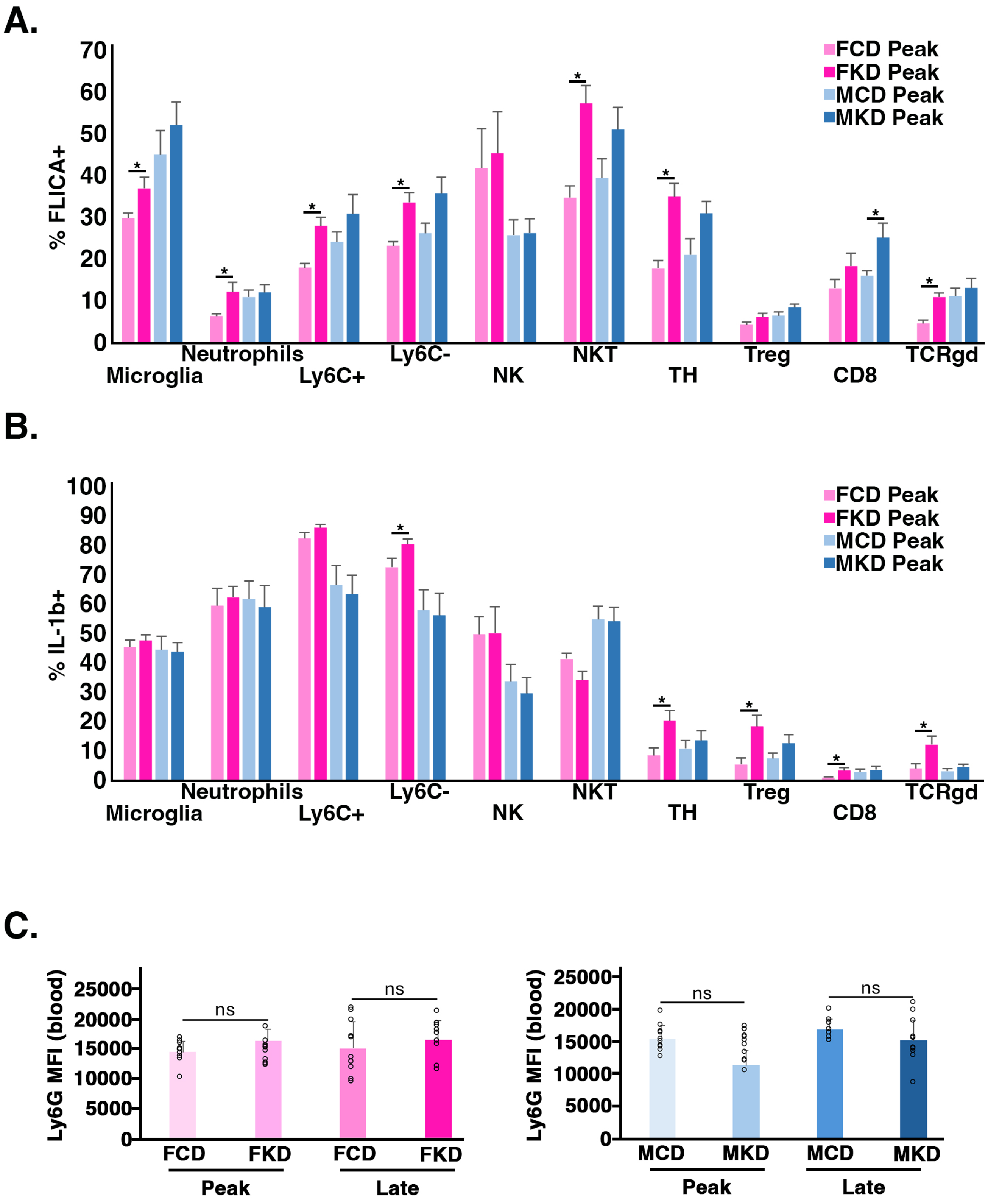

#### Supplemental Figure 5

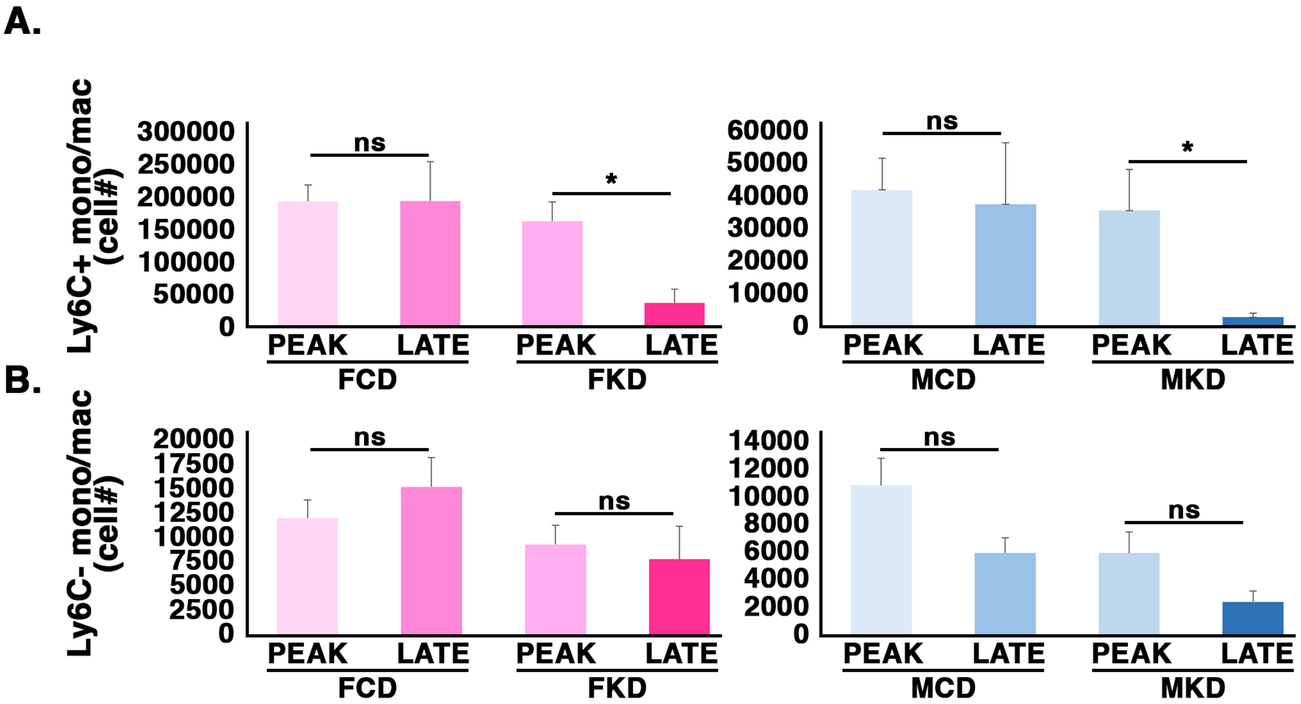

### Supplemental Figure 6

A.

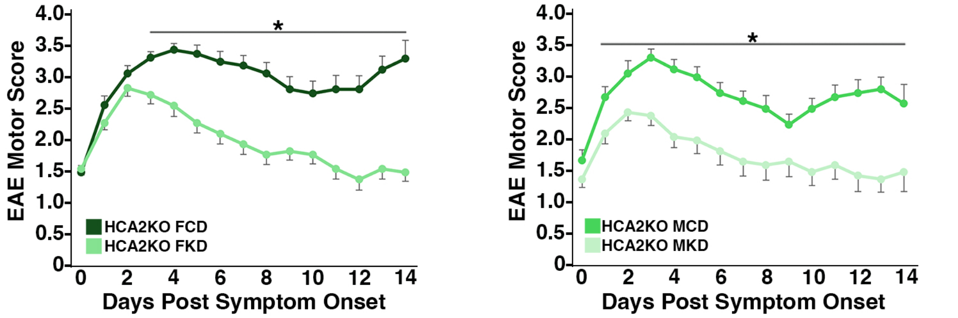


B.

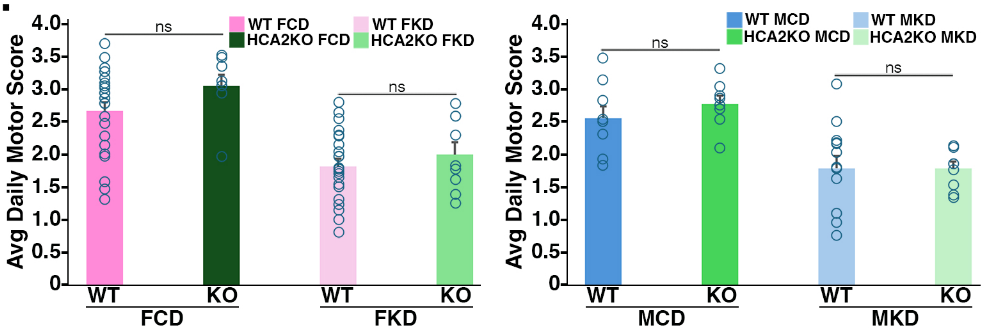


C.

| | FOLD CHNG v CD | |
|---|---|---|
| | SC | SC |
| | Late | Late |
| | HCA2KO | HCA2KO |
| | FKD | MKD |
| CD45hi | 0.37 | 0.16 |
| CD45+ | 0.42 | 0.26 |
| Bcell | | 0.32 |
| NK | 0.46 | 0.20 |
| NKT | 0.47 | 0.25 |
| CD11b+ | 0.35 | 0.09 |
| Neutr | 0.29 | 0.11 |
| Ly6C+ mono/mac | 0.43 | 0.06 |
| Ly6C- mono/mac | 0.28 | 0.19 |
| cDC | 0.27 | 0.06 |
| TH | 0.39 | 0.24 |
| Treg | 0.24 | 0.14 |
| CD8 | 0.43 | 0.51 |
| TCRgd | 0.87 | 0.05 |
| Microglia | | 0.67 |

D.

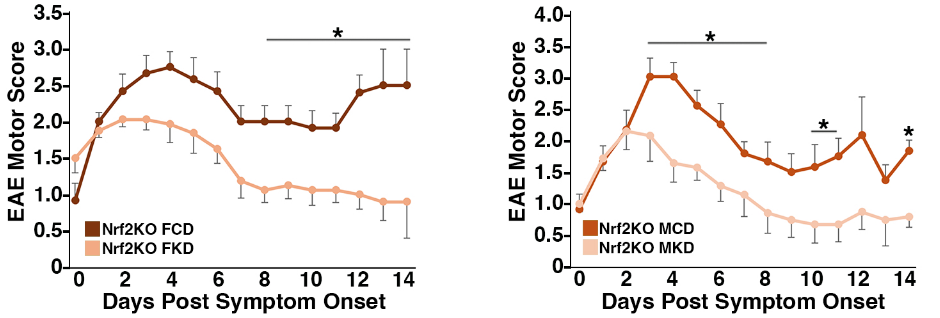


E.

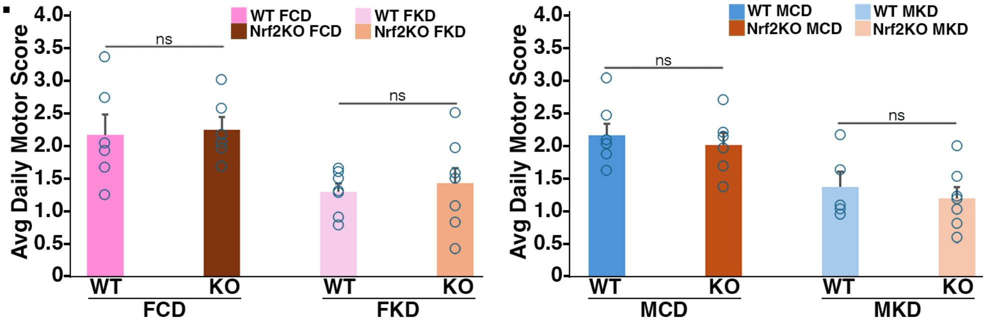

Supplemental Table 1

| Marker | Fluorochrome | Company | Catalog# |
|---|---|---|---|
| CD11b | APC Fire810 | Biolegend | 101287 |
| CD11c | Spark NIR685 | Biolegend | 117368 |
| CD19 | FITC | Biolegend | 152404 |
| CD19 | PE | Biolegend | 115507 |
| CD25 | BV785 | Biolegend | 102051 |
| CD3 | PE/Cy7 | Biolegend | 100220 |
| CD4 | AF700 | Biolegend | 100536 |
| CD44 | PerCP/Cy5.5 | Biolegend | 103032 |
| CD45.2 | Spark Blue574 | Biolegend | 285018 |
| CD62L | PEFire810 | Biolegend | 161205 |
| CD8a | APC Fire750 | Biolegend | 100765 |
| F4/80 | PE Dazzle594 | Biolegend | 123146 |
| IL-1b | PerCP-eFluor 710 | Invitrogen | 46-711-480 |
| Ly6C | BUV805 | BD Biosciences | 755202 |
| Ly6G | BUV395 | BD Biosciences | 563978 |
| MHCII | PE/Cy5 | Biolegend | 107611 |
| NK1.1 | FITC | Biolegend | 108706 |
| NK1.1 | PE | Biolegend | 108708 |
| NKp46 | BUV737 | BD Biosciences | 612805 |
| TCRβ | BV711 | Biolegend | 109243 |
| TCRγδ | BV650 | Biolegend | 118147 |

Supplemental Table 2

| NAME | ABBREVIATION | SURFACE MARKERS |
|---|---|---|
| **B Cell** | B Cell | CD45+ CD19+ |
| **Natural Killer** | NK | CD45+ NK1.1+ (or NKp46+) CD3- |
| **NK T Cell** | NKT | CD45+ NK1.1+ (or NKp46+) CD3+ |
| **Myeloid Cell** | CD11b+ | CD45+ CD19- CD11b+ |
| **Neutrophil** | Neutr | CD45+ CD19- CD11b+ $Ly6C^{int}$ Ly6G+ |
| **Monocyte/Macrophage** | Mono/Mac | CD45+ CD19- CD11b+ Ly6G- |
| **Ly6C- Monocyte/Macrophage** | Ly6C- | CD45+ CD19- CD11b+ Ly6G- Ly6C- |
| **Ly6C+ Monocyte/Macrophage** | Ly6C+ | CD45+ CD19- CD11b+ Ly6G- Ly6C+ |
| **Type 1 Classical Dendritic Cell** | cDC | CD45+ CD19- CD11b- CD3- CD11c+ |
| **T Helper Cell** | TH | CD45+ CD19- CD11b- CD3+ TCRβ+ CD4+ FoxP3- |
| **T Regulatory Cell** | Treg | CD45+ CD19- CD11b- CD3+ TCRβ+ CD4+ FoxP3+ |
| **CD8 Cytotoxic T Cell** | CD8 | CD45+ CD19- CD11b- CD3+ TCRβ+ CD8+ |
| **γδ T Cell** | TCRgd | CD45+ CD19- CD11b- CD3+ TCRγδ+ |
| **Microglia** | Microglia | $CD45^{lo}$ CD11b+ |
| **Monocytes** | Mono | $CD45^{hi}$ CD19- CD11b+ Ly6G- F4/80- |
| **Macrophage** | Mac | $CD45^{hi}$ CD19- CD11b+ Ly6G- F4/80+ |

### Supplemental Table 3 SC EAE

| COUNTS | avg SC PEAK FCD | avg SC PEAK FKD | sem SC PEAK FCD | sem SC PEAK FKD | FOLD CHNG SC PEAK FKD | Mann-Whit CD SC PEAK FKD | avg SC LATE FCD | avg SC LATE FKD | sem SC LATE FCD | sem SC LATE FKD | FOLD CHNG SC LATE FKD | Mann-Whit CD SC LATE FKD |
|---|---|---|---|---|---|---|---|---|---|---|---|---|
| CD45hi | 364862 | 271648 | 42856 | 42310 | 0.74 | 0.110 | 415095 | 93683 | 97098 | 37748 | **0.23** | ***0.003** |
| Bcells | 2440 | 1745 | 591 | 331 | 0.71 | 0.538 | 1280 | 480 | 426 | 161 | 0.37 | 0.086 |
| NK | 6708 | 4323 | 917 | 546 | 0.64 | 0.085 | 10720 | 3371 | 2619 | 925 | **0.31** | ***0.009** |
| NKT | 576 | 718 | 119 | 198 | 1.25 | 0.806 | 753 | 146 | 191 | 48 | **0.19** | ***0.002** |
| CD11b+ | 261683 | 185673 | 34204 | 32041 | 0.71 | 0.124 | 282673 | 49373 | 72999 | 25212 | **0.17** | ***0.002** |
| Neutr | 55834 | 13176 | 12673 | 2429 | **0.24** | ***0.006** | 73186 | 4531 | 13930 | 1469 | **0.06** | ***0.000** |
| Ly6C- mono/mac | 12081 | 9357 | 1883 | 1985 | 0.77 | 0.157 | 15308 | 7840 | 2997 | 3379 | **0.51** | ***0.022** |
| Ly6C+ mono/mac | 194065 | 163438 | 25450 | 29338 | 0.84 | 0.498 | 194433 | 37085 | 60932 | 21143 | **0.19** | ***0.004** |
| cDC | 12372 | 9926 | 1076 | 2285 | 0.80 | 0.110 | 15950 | 6425 | 3736 | 2997 | **0.40** | ***0.018** |
| TH | 43506 | 38104 | 4783 | 4629 | 0.88 | 0.424 | 47272 | 12673 | 11433 | 3100 | **0.27** | ***0.003** |
| Treg | 15214 | 11748 | 1816 | 1380 | 0.77 | 0.218 | 25394 | 7618 | 5348 | 2247 | **0.30** | ***0.006** |
| CD8 | 6666 | 6973 | 1018 | 1034 | 1.05 | 0.854 | 7010 | 3534 | 2172 | 1053 | 0.50 | 0.288 |
| TCRgd | 2691 | 2360 | 746 | 641 | 0.88 | 0.218 | 2917 | 1926 | 683 | 1392 | **0.66** | ***0.011** |
| Microglia | 45182 | 55176 | 3897 | 6865 | 1.22 | 0.295 | 84433 | 54005 | 12083 | 7585 | 0.64 | 0.121 |
| Mono | 33089 | 22053 | 5118 | 5161 | 0.67 | 0.140 | 50053 | 7917 | 12408 | 3503 | **0.16** | ***0.001** |
| Ly6C+Mono | 27735 | 16383 | 4464 | 3540 | 0.59 | 0.074 | 42752 | 3961 | 12059 | 1751 | **0.09** | ***0.001** |
| Ly6C-Mono | 5238 | 5564 | 1277 | 1714 | 1.06 | 0.667 | 7303 | 3949 | 1144 | 1887 | **0.54** | ***0.041** |
| Mac | 171923 | 150213 | 21917 | 26195 | 0.87 | 0.622 | 159388 | 36962 | 53737 | 20689 | **0.23** | ***0.007** |
| Ly6C+Mac | 159071 | 141870 | 21651 | 26074 | 0.89 | 0.667 | 149106 | 32399 | 52358 | 19062 | **0.22** | ***0.007** |
| Ly6C-Mac | 12734 | 8151 | 1674 | 1066 | 0.64 | 0.074 | 10197 | 4587 | 2619 | 1887 | **0.45** | ***0.027** |

| COUNTS | avg SC PEAK MCD | avg SC PEAK MKD | sem SC PEAK MCD | sem SC PEAK MKD | FOLD CHNG SC PEAK MKD | Mann-Whit CD SC PEAK MKD | avg SC LATE MCD | avg SC LATE MKD | sem SC LATE MCD | sem SC LATE MKD | FOLD CHNG SC LATE MKD | Mann-Whit CD SC LATE MKD |
|---|---|---|---|---|---|---|---|---|---|---|---|---|
| CD45hi | 175520 | 106961 | 38722 | 31439 | 0.61 | 0.178 | 119950 | 19826 | 37311 | 5064 | **0.17** | ***0.000** |
| Bcells | 2964 | 2451 | 1218 | 775 | 0.83 | 0.630 | 595 | 162 | 132 | 28 | **0.27** | ***0.001** |
| NK | 6165 | 4834 | 727 | 1258 | 0.78 | 0.630 | 10762 | 2388 | 2144 | 810 | **0.22** | ***0.001** |
| NKT | 1007 | 841 | 334 | 328 | 0.83 | 0.501 | 1193 | 375 | 283 | 131 | **0.31** | ***0.025** |
| CD11b+ | 102863 | 58500 | 30580 | 19589 | 0.57 | 0.211 | 61108 | 6239 | 27972 | 1924 | **0.10** | ***0.001** |
| Neutr | 49684 | 16685 | 24206 | 6700 | 0.34 | 0.564 | 17250 | 1121 | 8034 | 350 | **0.06** | ***0.001** |
| Ly6C- mono/mac | 10773 | 5889 | 1957 | 1523 | **0.55** | ***0.034** | 5907 | 2367 | 1089 | 779 | **0.40** | ***0.002** |
| Ly6C+ mono/mac | 42248 | 35852 | 9766 | 12772 | 0.85 | 0.501 | 37780 | 2734 | 19146 | 1235 | **0.07** | ***0.002** |
| cDC | 5995 | 3355 | 877 | 827 | 0.56 | 0.083 | 6957 | 978 | 1226 | 428 | **0.14** | ***0.000** |
| TH | 25280 | 15916 | 5544 | 4565 | 0.63 | 0.178 | 16602 | 4007 | 4130 | 823 | **0.24** | ***0.001** |
| Treg | 11982 | 6922 | 2240 | 1738 | 0.58 | 0.124 | 10289 | 1852 | 1625 | 482 | **0.18** | ***0.000** |
| CD8 | 5523 | 4193 | 891 | 983 | 0.76 | 0.386 | 3478 | 1549 | 488 | 199 | **0.45** | ***0.002** |
| TCRgd | 2467 | 1265 | 742 | 481 | 0.51 | 0.149 | 2176 | 601 | 1495 | 436 | **0.28** | ***0.001** |
| Microglia | 107074 | 61094 | 9593 | 5831 | 0.57 | 0.004 | 56398 | 30562 | 5699 | 4857 | **0.54** | ***0.005** |
| Mono | 11635 | 9090 | 2452 | 3078 | 0.78 | 0.211 | 9601 | 1083 | 3059 | 365 | **0.11** | ***0.000** |
| Ly6C+Mono | 9132 | 7708 | 2075 | 2712 | 0.84 | 0.336 | 6184 | 538 | 2560 | 199 | **0.09** | ***0.001** |
| Ly6C-Mono | 2474 | 1385 | 532 | 405 | 0.56 | 0.211 | 3350 | 541 | 502 | 172 | **0.16** | ***0.000** |
| Mac | 43182 | 34269 | 10843 | 11970 | 0.79 | 0.630 | 34242 | 4022 | 17186 | 1294 | **0.12** | ***0.000** |
| Ly6C+Mac | 35105 | 29797 | 8990 | 10809 | 0.85 | 0.564 | 31248 | 2144 | 16525 | 1024 | **0.07** | ***0.002** |
| Ly6C-Mac | 8103 | 4498 | 1932 | 1242 | 0.56 | 0.149 | 2934 | 1869 | 730 | 777 | **0.64** | ***0.015** |

### Supplemental Table 3 SC Healthy

| COUNTS | avg SC 2w fed Healthy FCD | avg SC 2w fed Healthy FKD | sem SC 2w fed Healthy FCD | sem SC 2w fed Healthy FKD | FOLD CHNG v CD SC 2w fed Healthy FKD | Mann-Whit CD SC 2w fed Healthy FKD |
|---|---|---|---|---|---|---|
| CD45hi | 3260 | 3613 | 611 | 745 | 1.11 | 0.749 |
| Bcells | 447 | 454 | 70 | 91 | 1.02 | 0.949 |
| NK | 141 | 123 | 22 | 16 | 0.88 | 0.482 |
| NKT | 30 | 35 | 4 | 7 | 1.18 | 0.565 |
| CD11b+ | 1891 | 2332 | 396 | 536 | 1.23 | 0.655 |
| Neutr | 1271 | 1624 | 283 | 393 | 1.28 | 0.482 |
| Ly6C- mono/mac | 160 | 207 | 24 | 40 | 1.29 | 0.406 |
| Ly6C+ mono/mac | 461 | 502 | 110 | 116 | 1.09 | 0.565 |
| cDC | 61 | 53 | 13 | 16 | 0.87 | 0.523 |
| TH | 65 | 66 | 19 | 24 | 1.01 | 0.798 |
| Treg | 3 | 4 | 1 | 1 | 1.22 | 0.655 |
| CD8 | 58 | 65 | 12 | 15 | 1.12 | 0.898 |
| TCRgd | 8 | 8 | 2 | 3 | 1.06 | 0.898 |
| Microglia | 16755 | 18406 | 1000 | 1063 | 1.10 | 0.180 |
| Mono | 183 | 155 | 51 | 40 | 0.85 | 0.848 |
| Ly6C+Mono | 155 | 127 | 48 | 37 | 0.82 | 0.798 |
| Ly6C-Mono | 28 | 29 | 5 | 5 | 1.04 | 0.848 |
| Mac | 436 | 550 | 80 | 114 | 1.26 | 0.523 |
| Ly6C+Mac | 289 | 349 | 65 | 80 | 1.21 | 0.565 |
| Ly6C-Mac | 147 | 201 | 19 | 42 | 1.37 | 0.482 |

| COUNTS | avg SC 2w fed Healthy MCD | avg SC 2w fed Healthy MKD | sem SC 2w fed Healthy MCD | sem SC 2w fed Healthy MKD | FOLD CHNG v CD SC 2w fed Healthy MKD | Mann-Whit CD SC 2w fed Healthy MKD |
|---|---|---|---|---|---|---|
| CD45hi | 3153 | 4351 | 1026 | 507 | 1.38 | 0.064 |
| Bcells | 518 | 427 | 131 | 32 | 0.82 | 0.817 |
| NK | 114 | 104 | 57 | 22 | 0.91 | 0.355 |
| NKT | 39 | 34 | 24 | 5 | 0.87 | 0.165 |
| CD11b+ | 1886 | 3258 | 710 | 472 | **1.73** | ***0.015** |
| Neutr | 1342 | 2288 | 544 | 428 | **1.70** | ***0.037** |
| Ly6C- mono/mac | 142 | 358 | 32 | 89 | **2.52** | ***0.037** |
| Ly6C+ mono/mac | 402 | 611 | 154 | 82 | **1.52** | ***0.037** |
| cDC | 46 | 46 | 25 | 10 | 1.00 | 0.247 |
| TH | 43 | 65 | 13 | 26 | 1.53 | 0.355 |
| Treg | 3 | 3 | 1 | 1 | 0.84 | 0.908 |
| CD8 | 56 | 72 | 10 | 17 | 1.30 | 0.728 |
| TCRgd | 5 | 6 | 1 | 2 | 1.19 | 0.954 |
| Microglia | 21882 | 26343 | 1805 | 765 | 1.20 | 0.064 |
| Mono | 146 | 328 | 56 | 58 | **2.25** | ***0.021** |
| Ly6C+Mono | 127 | 231 | 53 | 40 | 1.83 | 0.064 |
| Ly6C-Mono | 19 | 96 | 3 | 64 | 4.96 | 0.083 |
| Mac | 393 | 635 | 110 | 78 | **1.61** | ***0.037** |
| Ly6C+Mac | 263 | 358 | 100 | 63 | 1.36 | 0.056 |
| Ly6C-Mac | 131 | 277 | 29 | 82 | 2.12 | 0.132 |

### Supplemental Table 3 Blood EAE

| COUNTS | avg Blood PEAK FCD | avg Blood PEAK FKD | sem Blood PEAK FCD | sem Blood PEAK FKD | FOLD CHNG Blood PEAK FKD | ttest FCD Blood PEAK FKD | avg Blood LATE FCD | avg Blood LATE FKD | sem Blood LATE FCD | sem Blood LATE FKD | FOLD CHNG Blood LATE FKD | ttest FCD Blood LATE FKD |
|---|---|---|---|---|---|---|---|---|---|---|---|---|
| CD45+ | 410993 | 532609 | 31373 | 59836 | 1.30 | 0.097 | 466860 | 362112 | 53905 | 51923 | 0.78 | 0.177 |
| Bcells | 96726 | 94081 | 6943 | 10378 | 0.97 | 0.838 | 122751 | 57294 | 17523 | 9720 | **0.47** | ***0.004** |
| NK | 29125 | 19374 | 3334 | 2347 | **0.67** | ***0.025** | 29651 | 17495 | 3593 | 1216 | **0.59** | ***0.004** |
| NKT | 2353 | 2816 | 122 | 349 | 1.20 | 0.244 | 2048 | 1864 | 231 | 180 | 0.91 | 0.537 |
| CD11b+ | 206656 | 335394 | 28019 | 50968 | **1.62** | ***0.045** | 236660 | 239816 | 44360 | 45037 | 1.01 | 0.961 |
| Neutr | 141837 | 244984 | 20843 | 35988 | **1.73** | ***0.026** | 152734 | 196376 | 25168 | 39825 | 1.29 | 0.365 |
| Ly6C- mono | 17184 | 21741 | 1878 | 4687 | 1.27 | 0.395 | 19877 | 13658 | 4186 | 1537 | 0.69 | 0.178 |
| Ly6C+ mono | 43813 | 66343 | 7808 | 12601 | 1.51 | 0.154 | 58585 | 26556 | 17132 | 4961 | 0.45 | 0.088 |
| cDC | 1087 | 1325 | 125 | 421 | 1.22 | 0.608 | 1881 | 909 | 455 | 116 | 0.48 | 0.052 |
| TH | 33274 | 37718 | 2033 | 5640 | 1.13 | 0.485 | 27330 | 16670 | 3929 | 3683 | 0.61 | 0.062 |
| Treg | 2311 | 1700 | 260 | 181 | 0.74 | 0.065 | 1363 | 1091 | 214 | 162 | 0.80 | 0.321 |
| CD8 | 28498 | 31370 | 1710 | 3319 | 1.10 | 0.465 | 30354 | 20255 | 2802 | 1522 | **0.67** | ***0.005** |
| TCRgd | 1917 | 1640 | 175 | 253 | 0.86 | 0.389 | 1798 | 958 | 288 | 75 | **0.53** | ***0.010** |

| COUNTS | avg Blood PEAK MCD | avg Blood PEAK MKD | sem Blood PEAK MCD | sem Blood PEAK MKD | FOLD CHNG Blood PEAK MKD | ttest MCD Blood PEAK MKD | avg Blood LATE MCD | avg Blood LATE MKD | sem Blood LATE MCD | sem Blood LATE MKD | FOLD CHNG Blood LATE MKD | ttest MCD Blood LATE MKD |
|---|---|---|---|---|---|---|---|---|---|---|---|---|
| CD45+ | 370153 | 513302 | 59444 | 22651 | **1.39** | ***0.025** | 286133 | 504403 | 34802 | 67969 | **1.76** | ***0.021** |
| Bcells | 101415 | 130775 | 17618 | 11516 | 1.29 | 0.158 | 70584 | 55671 | 11532 | 6243 | 0.79 | 0.233 |
| NK | 25819 | 24417 | 4324 | 1917 | 0.95 | 0.753 | 17615 | 13451 | 1748 | 1101 | **0.76** | ***0.046** |
| NKT | 2863 | 3284 | 529 | 499 | 1.15 | 0.559 | 1115 | 1162 | 140 | 200 | 1.04 | 0.862 |
| CD11b+ | 172874 | 279521 | 27604 | 16052 | **1.62** | ***0.002** | 151159 | 385957 | 31889 | 61000 | **2.55** | ***0.007** |
| Neutr | 118272 | 198028 | 17342 | 12283 | **1.67** | ***0.001** | 121343 | 342065 | 28709 | 54434 | **2.82** | ***0.005** |
| Ly6C- mono | 13096 | 20545 | 2166 | 1819 | **1.57** | ***0.013** | 8577 | 13456 | 797 | 2275 | 1.57 | 0.100 |
| Ly6C+ mono | 38039 | 56922 | 8334 | 5556 | 1.50 | 0.062 | 19731 | 29409 | 2882 | 5843 | 1.49 | 0.209 |
| cDC | 1295 | 932 | 369 | 100 | 0.72 | 0.312 | 2909 | 1225 | 822 | 260 | **0.42** | ***0.035** |
| TH | 25423 | 32326 | 3657 | 2795 | 1.27 | 0.135 | 17407 | 21047 | 1912 | 1410 | 1.21 | 0.133 |
| Treg | 2061 | 2379 | 410 | 338 | 1.15 | 0.542 | 1275 | 1403 | 226 | 130 | 1.10 | 0.606 |
| CD8 | 27976 | 30368 | 4545 | 1781 | 1.09 | 0.602 | 17545 | 18793 | 2128 | 1007 | 1.07 | 0.565 |
| TCRgd | 1954 | 1390 | 393 | 105 | 0.71 | 0.145 | 1082 | 1420 | 121 | 145 | 1.31 | 0.110 |

### Supplemental Table 3 Blood Healthy

| COUNTS | avg BLOOD 2w fed Healthy FCD | avg BLOOD 2w fed Healthy FKD | sem BLOOD 2w fed Healthy FCD | sem BLOOD 2w fed Healthy FKD | FOLD CHNG v CD BLOOD 2w fed Healthy FKD | ttest CD BLOOD 2w fed Healthy FKD |
|---|---|---|---|---|---|---|
| CD45+ | 296767 | 241734 | 33437 | 14951 | 0.81 | 0.131 |
| Bcells | 107815 | 85671 | 17595 | 4169 | 0.79 | 0.212 |
| NK | 21085 | 13527 | 1783 | 1069 | **0.64** | ***0.002** |
| NKT | 3841 | 3300 | 506 | 864 | 0.86 | 0.573 |
| CD11b+ | 81263 | 73691 | 19574 | 10113 | 0.91 | 0.719 |
| Neutr | 50260 | 54538 | 16981 | 8207 | 1.09 | 0.812 |
| Ly6C- mono | 8456 | 6225 | 1003 | 556 | 0.74 | 0.056 |
| Ly6C+ mono | 19360 | 10216 | 4285 | 1340 | **0.53** | ***0.047** |
| cDC | 593 | 209 | 171 | 17 | **0.35** | ***0.031** |
| TH | 34072 | 28358 | 1390 | 2611 | 0.83 | 0.058 |
| Treg | 678 | 721 | 44 | 22 | 1.06 | 0.368 |
| CD8 | 34212 | 27555 | 2669 | 1297 | **0.81** | ***0.031** |
| TCRgd | 1265 | 799 | 162 | 47 | **0.63** | ***0.010** |

| COUNTS | avg BLOOD 2w fed Healthy MCD | avg BLOOD 2w fed Healthy MKD | sem BLOOD 2w fed Healthy MCD | sem BLOOD 2w fed Healthy MKD | FOLD CHNG v CD BLOOD 2w fed Healthy MKD | ttest CD BLOOD 2w fed Healthy MKD |
|---|---|---|---|---|---|---|
| CD45+ | 315431 | 505224 | 46595 | 38489 | **1.60** | ***0.006** |
| Bcells | 111832 | 111404 | 28962 | 15358 | 1.00 | 0.989 |
| NK | 22002 | 19069 | 3424 | 2381 | 0.87 | 0.474 |
| NKT | 2858 | 4592 | 670 | 2413 | 1.61 | 0.498 |
| CD11b+ | 101101 | 288958 | 15662 | 41714 | **2.86** | ***0.001** |
| Neutr | 71726 | 240391 | 11596 | 41725 | **3.35** | ***0.002** |
| Ly6C- mono | 7488 | 9188 | 1069 | 483 | 1.23 | 0.148 |
| Ly6C+ mono | 19390 | 33722 | 4857 | 4684 | **1.74** | ***0.045** |
| cDC | 455 | 464 | 115 | 96 | 1.02 | 0.952 |
| TH | 33113 | 31983 | 5773 | 3595 | 0.97 | 0.864 |
| Treg | 871 | 1090 | 148 | 99 | 1.25 | 0.218 |
| CD8 | 28052 | 27269 | 5789 | 2563 | 0.97 | 0.898 |
| TCRgd | 802 | 930 | 155 | 73 | 1.16 | 0.443 |

### Supplemental Table 3 BM EAE

| COUNTS | avg BM PEAK FCD | avg BM PEAK FKD | sem BM PEAK FCD | sem BM PEAK FKD | FOLD CHNG BM PEAK FKD | ttest CD BM PEAK FKD | avg BM LATE FCD | avg BM LATE FKD | sem BM LATE FCD | sem BM LATE FKD | FOLD CHNG BM LATE FKD | ttest CD BM LATE FKD |
|---|---|---|---|---|---|---|---|---|---|---|---|---|
| CD45+ | 29317400 | 20788614 | 1156899 | 1512068 | **0.71** | ***0.000** | 27700727 | 26541795 | 2101702 | 2192186 | 0.96 | 0.707 |
| Bcells | 1112495 | 862851 | 195390 | 142419 | 0.78 | 0.309 | 2046155 | 1593627 | 288447 | 261781 | 0.78 | 0.259 |
| NK | 230803 | 148216 | 17879 | 11351 | **0.64** | ***0.001** | 207220 | 220679 | 16381 | 17390 | 1.06 | 0.579 |
| NKT | 225088 | 182290 | 22470 | 11443 | 0.81 | 0.097 | 222703 | 223875 | 16340 | 17085 | 1.01 | 0.961 |
| CD11b+ | 25677782 | 18227237 | 1099107 | 1418098 | **0.71** | ***0.001** | 22472128 | 22497251 | 1907401 | 2104876 | 1.00 | 0.993 |
| Neutr | 19392715 | 13783575 | 824155 | 1197320 | **0.71** | ***0.001** | 16284504 | 16875842 | 1459918 | 1876629 | 1.04 | 0.806 |
| Ly6C- mono | 165146 | 128449 | 14195 | 11043 | 0.78 | 0.053 | 168834 | 192067 | 13815 | 14583 | 1.14 | 0.261 |
| Ly6C+ mono | 5460011 | 3873869 | 378173 | 285258 | **0.71** | ***0.003** | 5988480 | 5274280 | 473790 | 357847 | 0.88 | 0.243 |
| cDC | 309406 | 171628 | 28378 | 14311 | **0.55** | ***0.000** | 419613 | 277498 | 42584 | 36327 | **0.66** | ***0.020** |
| TH | 49291 | 41571 | 10735 | 10717 | 0.84 | 0.617 | 52479 | 49062 | 12063 | 7288 | 0.93 | 0.811 |
| Treg | 55247 | 35321 | 9080 | 6135 | 0.64 | 0.080 | 47256 | 43818 | 9556 | 6348 | 0.93 | 0.768 |
| CD8 | 88592 | 78808 | 17672 | 16965 | 0.89 | 0.694 | 111044 | 104357 | 23019 | 14754 | 0.94 | 0.809 |
| TCRgd | 15476 | 11124 | 2155 | 1658 | 0.72 | 0.122 | 18115 | 15244 | 2428 | 1400 | 0.84 | 0.318 |

| COUNTS | avg BM PEAK MCD | avg BM PEAK MKD | sem BM PEAK MCD | sem BM PEAK MKD | FOLD CHNG BM PEAK MKD | ttest CD BM PEAK MKD | avg BM LATE MCD | avg BM LATE MKD | sem BM LATE MCD | sem BM LATE MKD | FOLD CHNG BM LATE MKD | ttest CD BM LATE MKD |
|---|---|---|---|---|---|---|---|---|---|---|---|---|
| CD45+ | 24920250 | 26641773 | 1245786 | 1488441 | 1.07 | 0.386 | 32677796 | 32794902 | 1918500 | 1476290 | 1.00 | 0.961 |
| Bcells | 1395925 | 1187658 | 157986 | 120494 | 0.85 | 0.307 | 3019516 | 1416972 | 353205 | 271524 | **0.47** | ***0.002** |
| NK | 300015 | 310751 | 23244 | 33189 | 1.04 | 0.794 | 318838 | 209115 | 27142 | 18075 | **0.66** | ***0.002** |
| NKT | 140882 | 125246 | 16926 | 18523 | 0.89 | 0.540 | 189034 | 160124 | 17583 | 15322 | 0.85 | 0.233 |
| CD11b+ | 20708206 | 22797641 | 1111892 | 1341967 | 1.10 | 0.245 | 25836843 | 28911291 | 1440809 | 1413883 | 1.12 | 0.155 |
| Neutr | 15595136 | 17413234 | 873253 | 966794 | 1.12 | 0.178 | 19687477 | 21731995 | 1118701 | 1539762 | 1.10 | 0.337 |
| Ly6C- mono | 167965 | 224010 | 14579 | 30590 | 1.33 | 0.114 | 158642 | 157176 | 24259 | 8197 | 0.99 | 0.948 |
| Ly6C+ mono | 4563187 | 4727581 | 252263 | 365121 | 1.04 | 0.715 | 5604367 | 6380835 | 359270 | 499087 | 1.14 | 0.262 |
| cDC | 290649 | 312442 | 18886 | 30468 | 1.07 | 0.550 | 426665 | 249370 | 48594 | 27054 | **0.58** | ***0.003** |
| TH | 50846 | 39813 | 7063 | 3776 | 0.78 | 0.184 | 56117 | 30112 | 7077 | 3883 | **0.54** | ***0.002** |
| Treg | 60639 | 52860 | 5059 | 3999 | 0.87 | 0.242 | 75505 | 37598 | 11998 | 4693 | **0.50** | ***0.003** |
| CD8 | 108091 | 91744 | 11480 | 8040 | 0.85 | 0.257 | 146174 | 75655 | 16559 | 13883 | **0.52** | ***0.004** |
| TCRgd | 21720 | 15998 | 2411 | 1300 | **0.74** | ***0.050** | 24299 | 13258 | 2152 | 1529 | **0.55** | ***0.000** |

### Supplemental Table 3 BM Healthy

| COUNTS | avg BM 2w fed Healthy FCD | avg BM 2w fed Healthy FKD | sem BM 2w fed Healthy FCD | sem BM 2w fed Healthy FKD | FOLD CHNG v CD BM 2w fed Healthy FKD | ttest CD BM 2w fed Healthy FKD |
|---|---|---|---|---|---|---|
| CD45+ | 17318875 | 19349156 | 2496492 | 1404522 | 1.12 | 0.461 |
| Bcells | 1549707 | 1806710 | 288020 | 146794 | 1.17 | 0.410 |
| NK | 149768 | 175447 | 30657 | 16149 | 1.17 | 0.441 |
| NKT | 179538 | 178169 | 28274 | 11997 | 0.99 | 0.963 |
| CD11b+ | 12367733 | 13979954 | 1970735 | 685681 | 1.13 | 0.423 |
| Neutr | 8518758 | 9914097 | 1387769 | 451197 | 1.16 | 0.324 |
| Ly6C- mono | 100511 | 91438 | 14731 | 8580 | 0.91 | 0.578 |
| Ly6C+ mono | 3308335 | 3442107 | 552709 | 208121 | 1.04 | 0.812 |
| cDC | 312548 | 227561 | 33014 | 38556 | 0.73 | 0.095 |
| TH | 57254 | 74580 | 14531 | 11533 | 1.30 | 0.335 |
| Treg | 48387 | 55936 | 13485 | 10459 | 1.16 | 0.644 |
| CD8 | 148278 | 212395 | 47601 | 42486 | 1.43 | 0.301 |
| TCRgd | 28058 | 29968 | 5329 | 4367 | 1.07 | 0.771 |

| COUNTS | avg BM 2w fed Healthy MCD | avg BM 2w fed Healthy MKD | sem BM 2w fed Healthy MCD | sem BM 2w fed Healthy MKD | FOLD CHNG v CD BM 2w fed Healthy MKD | ttest CD BM 2w fed Healthy MKD |
|---|---|---|---|---|---|---|
| CD45+ | 19318688 | 24090019 | 1482040 | 605253 | **1.25** | ***0.007** |
| Bcells | 1789117 | 973499 | 121614 | 119141 | **0.54** | ***0.000** |
| NK | 183746 | 185454 | 24129 | 11547 | 1.01 | 0.947 |
| NKT | 127702 | 128005 | 10562 | 7639 | 1.00 | 0.981 |
| CD11b+ | 14043398 | 20683968 | 1150051 | 654325 | **1.47** | ***0.000** |
| Neutr | 10246610 | 15701782 | 704327 | 438222 | **1.53** | ***0.000** |
| Ly6C- mono | 107735 | 98661 | 9558 | 7226 | 0.92 | 0.443 |
| Ly6C+ mono | 3196915 | 4307366 | 398817 | 252299 | **1.35** | ***0.027** |
| cDC | 262807 | 208156 | 13359 | 15644 | **0.79** | ***0.016** |
| TH | 41970 | 19935 | 7614 | 1791 | **0.47** | ***0.010** |
| Treg | 31987 | 17985 | 3991 | 3580 | **0.56** | ***0.017** |
| CD8 | 85798 | 42557 | 16806 | 4973 | **0.50** | ***0.020** |
| TCRgd | 19629 | 10542 | 1806 | 942 | **0.54** | ***0.000** |

### Supplemental Table 4

#### 2 weeks of KD or CD feeding

| | FOLD CHNG v CD** | | FOLD CHNG v CD** | |
|---|---|---|---|---|
| | Blood | Blood | BM | BM |
| | Healthy | Healthy | Healthy | Healthy |
| | FKD | MKD | FKD | MKD |
| CD45+ | | 1.60 | | 1.25 |
| Bcells | | | | 0.54 |
| NK | 0.64 | | | |
| NKT | | | | |
| CD11b+ | | 2.86 | | 1.47 |
| Neutr | | 3.35 | | 1.53 |
| Ly6C- mono/mac | 0.72 | | | |
| Ly6C+ mono/mac | 0.53 | 1.74 | | 1.35 |
| cDC | 0.35 | | | 0.79 |
| TH | | | | 0.47 |
| Treg | | | | 0.56 |
| CD8 | 0.81 | | | 0.50 |
| TCRgd | 0.63 | | | 0.54 |

| KD/CD FOLD CHNG |
|---|
| 3.00 |
| 2.00 |
| 1.50 |
| 1.00 |
| 0.75 |
| 0.50 |
| 0.25 |
| 0.00 |
